# Variation of female pronucleus reveal oocyte or embryo abnormality: An expert experience deep learning of non-dark box analysis

**DOI:** 10.1101/2021.12.17.473071

**Authors:** Jingwei Yang, Yikang Wang, Chong Li, Wei Han, Weiwei Liu, Shun Xiong, Qi Zhang, Keya Tong, Guoning Huang, Xiaodong Zhang

## Abstract

**Background:** Pronuclear assessment appears to have the ability to distinguish good and bad embryos in the zygote stage, but paradoxical results were obtained in clinical studies. This situation might be caused by the robust qualitative detection of the development of dynamic pronuclei. Here, we aim to establish a quantitative pronuclear measurement method by applying expert experience deep learning from large annotated datasets.

**Methods:** Convinced handle-annotated 2PN images (13419) were used for deep learning then corresponded errors were recorded through handle check for subsequent parameters adjusting. We used 790 embryos with 52479 PN images from 155 patients for analysis the area of pronuclei and the pre-implantation genetic test results. Establishment of the exponential fitting equation and the key coefficient β 1was extracted from the model for quantitative analysis for pronuclear(PN) annotation and automatic recognition.

**Findings:** Based on the female original PN coefficient β1, the chromosome-normal rate in the blastocyst with biggest PN area is much higher than that of the blastocyst with smallest PN area (58.06% vs. 45.16%, OR=1.68 [1.07–2.64]; *P*=0.031). After adjusting coefficient β1 by the first three frames which high variance of outlier PN areas was removed, coefficient β1 at 12 hours and at 14 hours post-insemination, similar but stronger evidence was obtained. All these discrepancies resulted from the female propositus in the PGT-SR subgroup and smaller chromosomal errors.

**Conclusion(s):** The results suggest that detailed analysis of the images of embryos could improve our understanding of developmental biology.

**Funding:** None

## Introduction

Human embryos begin with the fertilization of an oocyte by a spermatozoon. A spermatozoon penetrates into the oocyte, causing a series of events that can be observed through a microscope, such as the cortical granule reaction, which prevents poly-fertilization, extrusion of the second polar body, and the formation and migration of two separate pronuclei that contain maternal and paternal chromosomes, respectively. The male and female pronuclei form in proximity of the zygote’s surface. Then, they need to move inwards in order to unite the paternal and maternal chromosomes on the first mitotic spindle (Scheffler K et al., 2021). During pronucleus (PN) migration from the periphery inward to the center of the zygote, the areas of both pronuclei increase gradually.

Phenomena related to pronuclear and nucleolar movements were first described by Wright et al.(Wright G et al., 1990). Notions including pronuclear alignment, and uneven/even numbers of chromosomes in the pronucleus and nucleolus precursor bodies (NPBs) have been expressed in more distinct pronuclear scores and used as a means to select embryos based on the Z-score (Scott LA et al., 1998). The scores have been correlated with improved embryo development (Balaban B et al., 2001; Rienzi L et al., 2002), increased pregnancy and implantation (Tesarik J et al.,2000; Zollner U et al.,2002;Jaroudi K et al., 2004), and embryonic chromosomal content (Gianaroli L et al., 2007; Gámiz P et al., 2003; Roos Kulmann MI et al., 2020) after the pre-implantation genetic test (PGT). However, some studies have disputed the effect of pronuclear scores for in vitro fertilization (IVF) or intracytoplasmic sperm injection (ICSI) (Nicoli A et al., 2013; Aydin S et al., 2011; Bar-Yoseph H et al., 2011), even if 0PN- and 1PN-derived blastocysts have similar neonatal results as 2PN-derived blastocysts (Doody KJ. 2021; Li M et al., 2021)

In theory, the pronuclear stage could be the only way to mirror the internal quality of the chromosomal integrity of the oocyte and the spermatozoon (Kuliev A et al., 2011;Lamb NE et al., 1996; Roos Kulmann MI et al., 2020). Meanwhile, developmental details such as disorder cleavage, embryonic fragment extrusion, uneven blastomeres, and abnormal morphokinetics during the post-zygote stage (cleavage, morula, and blastocyst stage) might reflect embryonic developmental dysfunction, mainly aneuploidy and mosaicism (Alpha Scientists in Reproductive M, 2011; Coticchio G et al., 2018; Munné S, 2006; Daughtry BL et al., 2019; Chavez SL et al., 2012). Due to the vague standard of methods (more than 6 scoring systems) in current pronuclear assessments (Nicoli A et al., 2013), the effect of pronuclear scores remains unclear. The dynamic character of the pronucleus, incongruent practice in IVF laboratories such as fertilization time and checking time, and the heterogeneity in patients make efficient qualitative classification for pronuclear assessment impossible.

Here, we aim to construct a computer-assisted algorithm for quantitative analysis for pronuclear assessment in ICSI patients from time-lapse incubators and test its efficacy in the diagnosis of chromosomal integrity in oocytes or embryos.

## Methods and materials

The deep learning images were obtained from the intracytoplasmic sperm injection (ICSI) cycles of 184 infertile couples requiring assisted reproductive technology (ART) therapy performed in 2019–2020 at the Reproductive and Genetic Institute of Chongqing in China. Infertility was diagnosed according to either female or male chromosomal/genomic abnormality (pre-implantation genetic test for chromosomal structural rearrangements [PGT-SR]), spontaneous abortion history (pre-implantation genetic test for aneuploidies [PGT-A]), unexplained reason (PGT-A), and tubal and pelvic factors combined with male chromosomal/genomic abnormality for ICSI and subsequent PGT-A. The study was approved by the local ethics committee. In total, 155 couples with 790 blastocyst-stage embryos were included in the final analysis, with 412 and 378 embryos in the PGT-A and PGT-SR subgroups, respectively (Figure 1).

**Fig.1.**
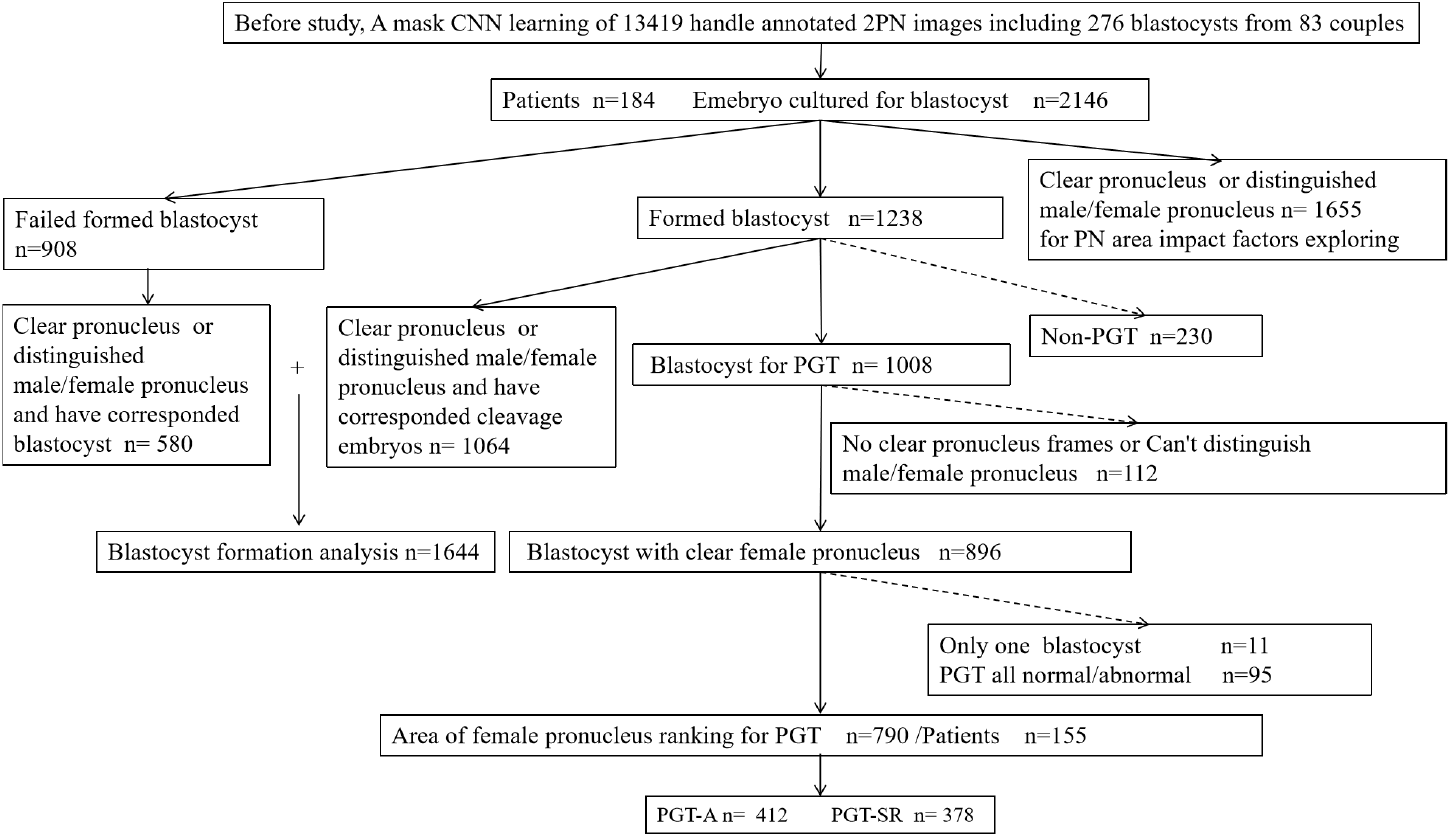
Flow chart

### Practices in ART

Before the ovaries were stimulated with recombinant FSH (Gonal-F, Merck Serono, Switzerland), downregulation was performed using a GnRH agonist (Decapeptyl; Ferring, Switzerland). Next, hCG (Ovidrel; Merck Serono, Italy) was administered when at least three leading follicles attained a mean diameter of >18 mm.

The flexible GnRH antagonist regimen included rFSH (Gonal-F; Serono, Aubonne, Switzerland) injection starting on day 2 of the menstrual cycle. The starting dose of rFSH was 75–300 IU daily and was customized according to the patient’s age, body mass index, antral follicle count, and baseline E2, P, FSH, and LH concentrations. Cetrorelix acetate (Cetrotide; Merck Serono Ltd., Aubonne, Switzerland) was used as the GnRH antagonist. Treatment with rFSH and cetrorelix acetate was continued until the day of the final oocyte maturation trigger.

Transvaginal oocyte retrieval was performed 36h after hCG injection. Cumulus-enclosed oocytes were collected in 2.5 ml of IVF medium (G-IVF, Vitrolife Sweden AB, Sweden) and incubated at 37°C under 5% O_2_ and 6% CO_2_ conditions for insemination.

Furthermore, sperm cells with normal morphology were selected, immobilized, and then microinjected into the oocyte cytoplasm 2–4 h after oocyte retrieval. Injected oocytes were then transferred into G-1 (Vitrolife, Sweden) medium droplets and placed into microwells of a custom-made well-of-the-well dish (EmbryoSlide®, Vitrolife Sweden AB, Sweden) containing 50 μl of equilibrated G-1 (Vitrolife Sweden AB, Sweden) microdroplets over the microwells and covered with 2.5 ml of Ovoil (Vitrolife Sweden AB, Sweden). Subsequently, the dish was immediately stored in a time-lapse (TL) system (EmbryoScope™, Vitrolife, Göteborg, Sweden). After 3 days of culture, the embryos were extracted and transferred to a new well-of-the-well dish containing 50 μl of equilibrated G-2 (Vitrolife Sweden AB, Sweden) microdroplets over the microwells and covered with 2.5 ml of Ovoil (Vitrolife Sweden AB, Sweden). The TL image acquisition was set every 10–15 min at seven different focal planes for each embryo. Images (1280 × 1024 pixels) were acquired using a Leica 20 × 0.40 LWD Hoffman Modulation contrast objective specialized for 635-nm illumination.

Transferable blastocysts were defined as follows: at least in the blastocyst stage at day 5 (120 h after ICSI) with moderate expansion, having easily discernible tightly compacted inner cell mass (ICM), and having trophectoderm (TE) either in many cells forming a cohesive epithelium or in few cells forming a loose epithelium.

### TL setting

The data are multi-view Hoffmann modulation contrast (HMC) microscopic images of developing cells in 11 different focal segments (−75, −60, −45, −30, −15, 0, 15, 30, 45, 60, 75) taken every 15 minutes. HMC is a kind of oblique lighting technology commonly used in IVF (Hoffman R et al,.1975). When oblique light irradiates the sample, it refracts and diffracts. The light line generates different shadows through the objective lens optical density regulator, so that the surface of the transparent sample produces a light and shade difference in order to enhance the contrast. The diameter of EmbryoSlide® (Vitrolife, Switzerland) is 250 μm. Therefore, the total area of the well was 49062.5 μm^2^. We measured the number of pixels of the well of the culture dish in all the time-lapse images. The number of pixels inside the well was 16077.98 ± 192.35. The relationship between a pixel and its actual size was 1 pixel = 0.3275 μm^2^(Zhao M et al,.2021).

### Establishment of the algorithm for quantitative analysis for pronuclear assessment

#### (i) Pronuclear annotation and automatic recognition and labeling

For the accurate measuring of PN edge and area, a pre-processing of TL images, a Laplacian-based method that could confirm the clearest focal plane from the 11 Z-stack images(Cai D et al,.2006; Belkin, M et al,.2005) was employed (Figure 2A). Then, expert experience features training for recognizing perivitelline space and PN was performed by a mask region-convolutional neural network (Mask R-CNN), which allowed us to easily estimate PN poses in the same framework (He K et al,.2020). An abandoned mechanism was introduced to automatic pronucleus recognition, and any abnormal images (0, 1, 3, or more than 3PN) were discarded.

**Fig.2.**
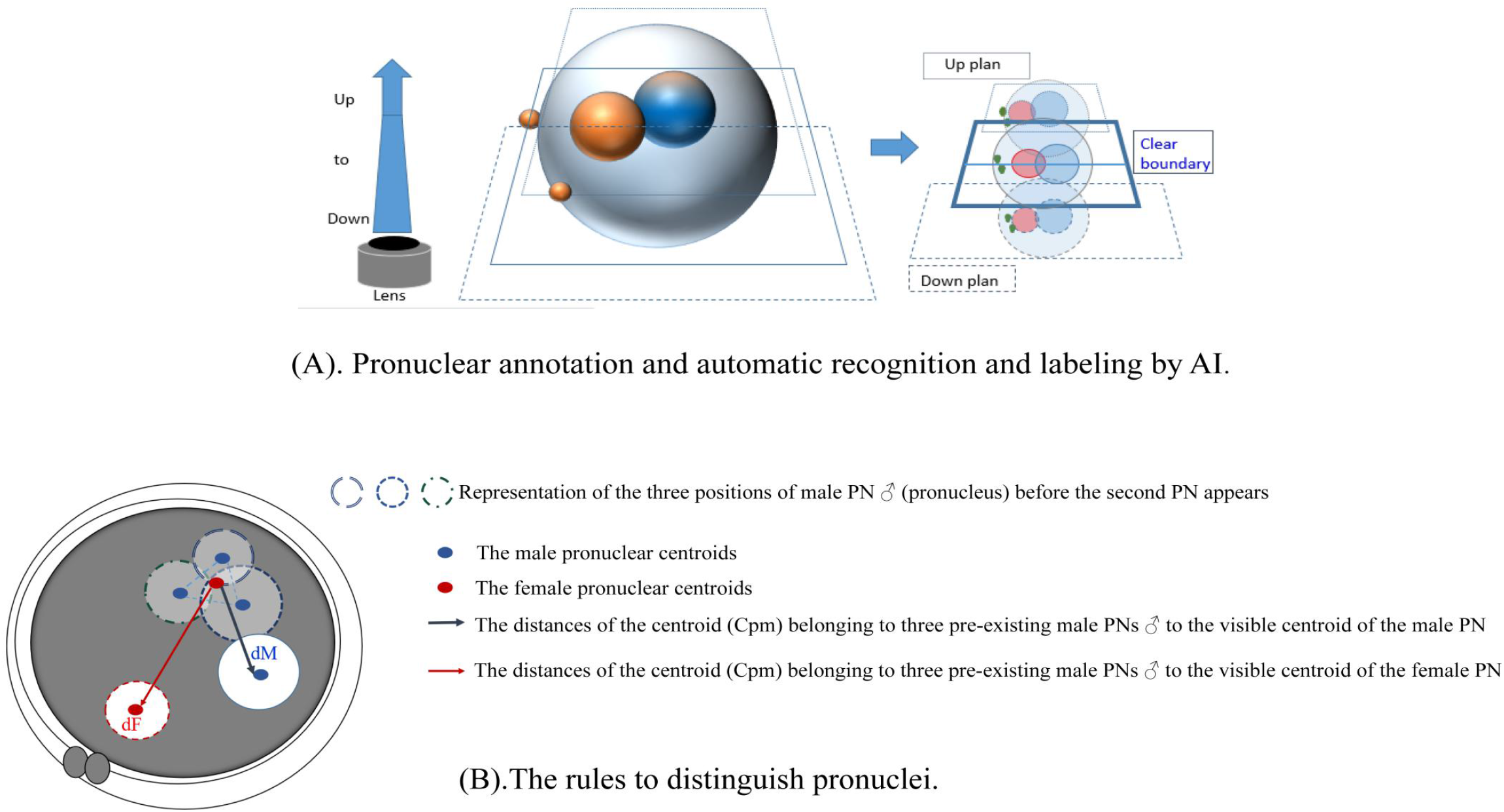
AI automatic recognition in PNs

**Fig.2.**
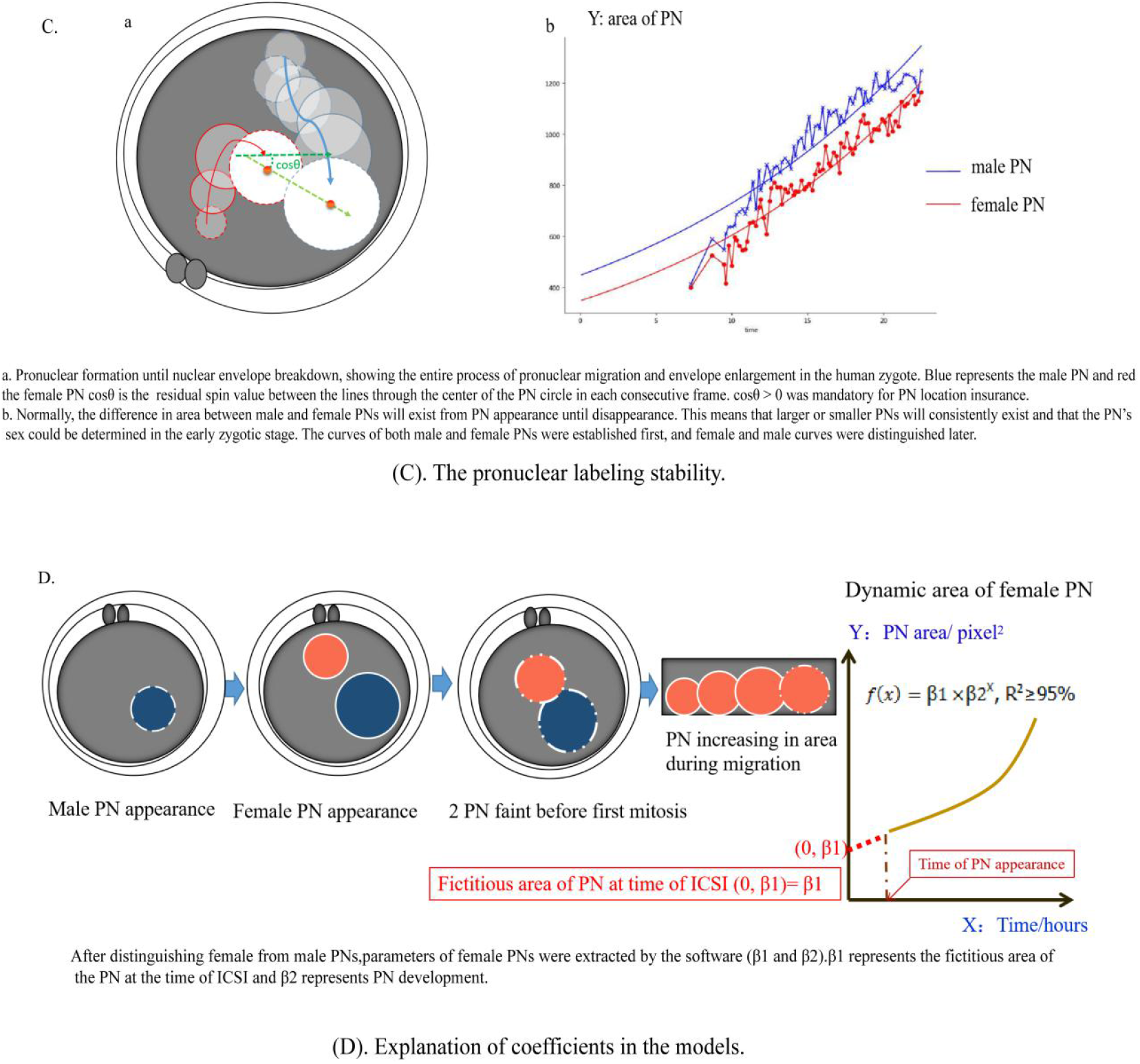
AI automatic recognition in PNs

### (ii) Distinguishing female and male pronuclei

Normally, two separate pronuclei appear at different times and positions inside the perivitelline space. Three main approaches can be employed for PN classification. First, the position of two separate pronuclei can be considered. The female pronucleus is closer to the second polar body (PB) than the male pronucleus. Second, the male pronucleus appears earlier than the female pronucleus, but the sequence of pronuclear appearance is hard to differentiate sometimes due to the image quality. Third, male pronuclei are larger than female pronuclei in the early zygote stage (Wiker S et al,.1990). However, due to the inherent limitations in automatic labeling, potential inaccurate labeling will be ignored in data outputting; the simpler the annotation, the higher the efficacy that might be obtained in practice. Thus, the PB was not employed as a feature for machine learning but for following handle checking and correction (Figure 3A, type and proportion). Only the second and third methods were employed to distinguish pronuclei and for automatic PN identification, and the second method was employed prior to the third method for automatic PN identification (Figure 2B). All pronuclear identifications by computer were confirmed and corrected by a senior embryologist, who did not know the PGT results.

**Fig.3.**
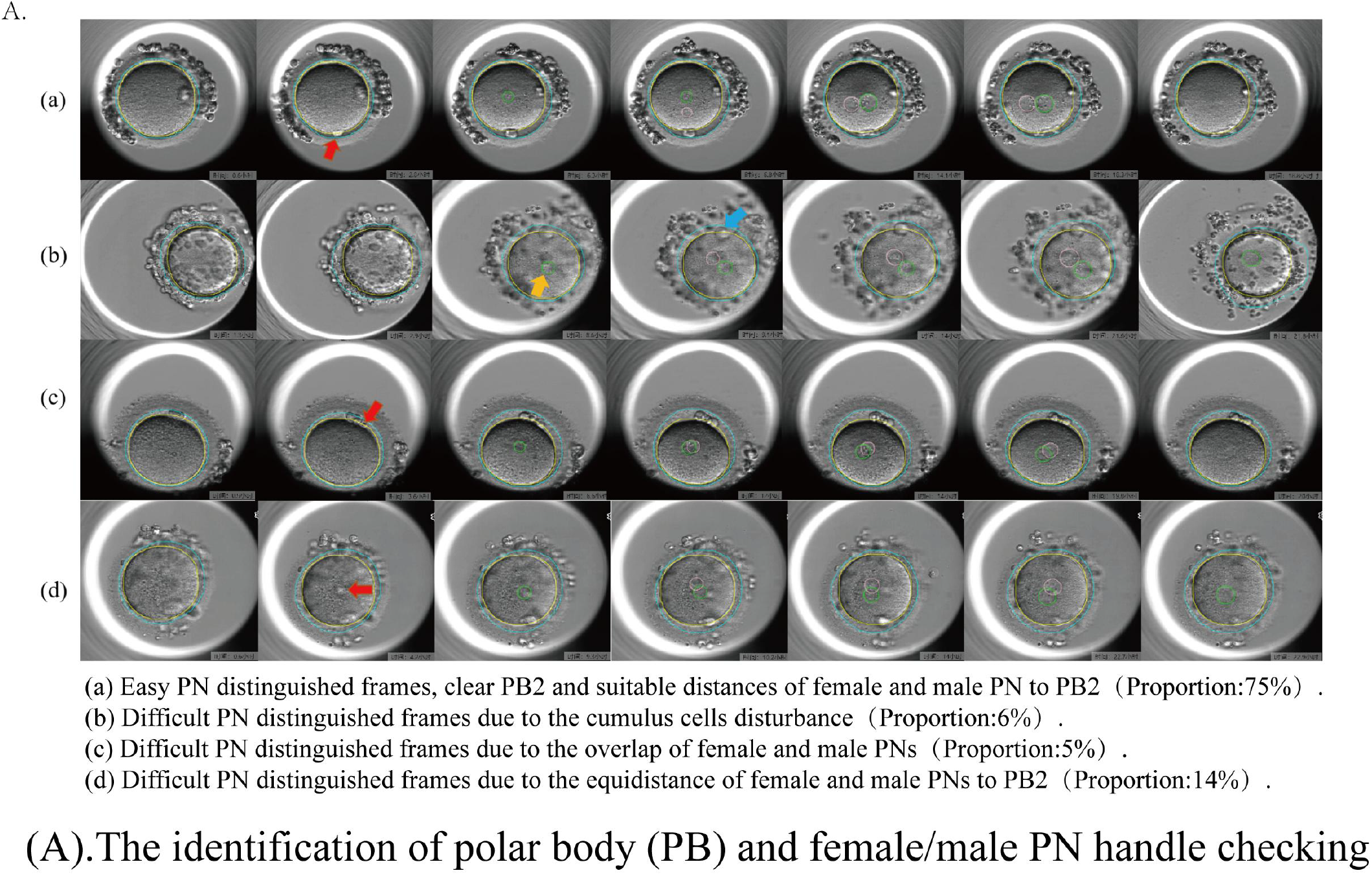
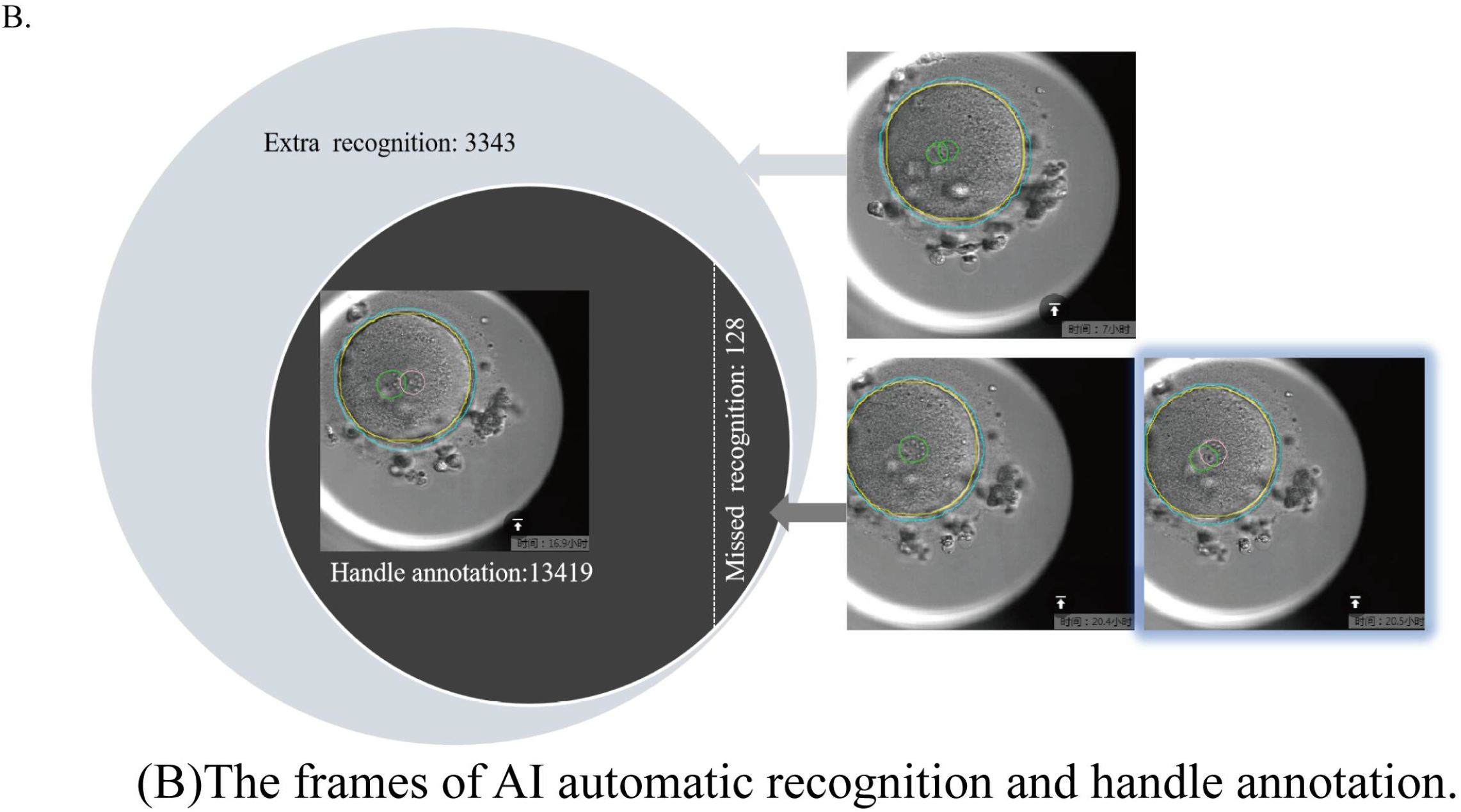
The statement of PN develpmental model and adjustment of coefficient

### (iii) The pronuclear labeling stability

In practice, after the orientation of female and male pronuclei, AI might produce errors when separately labeling and categorizing female or male PNs, so vector calculus discrimination through cosine similarity (cosθ > 0) was performed for PN location insurance in subsequence images (Figure 2C-a).

#### (iv) Exponential model for coefficient extraction

Mathematical models were employed to describe the dynamic nature of pronuclear development, including linear, logarithmic, cosine, quadratic, and exponential functions. Finally, an exponential fitting equation (Figure 2C-b) *f* (*x*) = β1 ×β2^x^ was employed and the key coefficient β1 was extracted from the model (for all others, R^2^ < 90%). The fitting degree (coefficient of correlation, R^2^) in the exponential model ranged from 98% to 99.99%.

#### (v) Explanation of coefficients in mathematical models

The high value of R^2^ implies that the development of the PN (from appearance to disappearance) complies with the exponential mathematical model. β1 represents the fictitious area of the PN at the time of ICSI (*f* (0) = β1 × β2^0^, where (*f* (*time of ICSI t* = 0 = β1 ×1 and β2^0^= 1) and β2 represents the PN development trend (Figure 2D). From this model, any value of the PN area (from 6 to 22 hours after ICSI) could be obtained. However, because β2 was approximately 1 (original β2 mean ± SD: 1.04 ± 0.017, range 1.01 to 1.11), the object of the study was β1.

### Chromosomal detection in blastocyst-stage embryos

On day 5 (120 h after ICSI), embryos with visible blastocoele were considered as blastocysts without taking quality in consideration.

Transferable blastocysts were defined as follows: at least in the blastocyst stage at day 5 with moderate expansion, having easily discernible tightly compacted inner cell mass (ICM) and having trophectoderm (TE) either in many cells forming a cohesive epithelium or in few cells forming a loose epithelium.

Several TE cells were extracted for biopsy using mechanical blunt dissection (Yang D et al., 2020). Following biopsy, the cells were placed into 0.2-mL thinly walled tubes, which were sealed and frozen by placing them in a freezer at −20°C prior to genetic screening. Single-cell, whole-genome amplification (WGA) with multiple annealing and looping-based amplification cycles (MALBAC) was used.

We performed WGA on cleavage-stage blastomeres using MALBAC following the manufacturer’s protocol (Catalog No. YK001B; Yikon Genomics).

Cells were lysed by heating (20 min at 50°C and 10 min at 80°C) in 5 μL of lysis buffer. Then, 30 μL of freshly prepared pre-amplification mix was added to each tube and the mixture was incubated at 94°C for 3 min. Next, DNA was amplified using 8 cycles of 40 s at 20°C, 40 s at 30°C, 30 s at 40°C, 30 s at 50°C, 30 s at 60°C, 4 min at 70°C, 20 s at 95°C, and 10 s at 58°C and immediately placed on ice. We then added 30 μL of the amplification reaction mix to each tube and incubated the mixture at 94°C for 30 s, followed by 17 cycles of 20 s at 94°C, 30 s at 58°C, and 3 min at 72°C.

Low-coverage (0.3×), genome-sequenced MALBAC products were purified using a DNA purification kit to construct our DNA library.

### Chromosomal error definition

“Chromosomal-normal” was defined as follows: subsequently developed embryos (blastocysts) were euploid, without genomic disorders of deletions and/or duplications (including microdeletions and/or -duplications), i.e., 46 XN.

“Sole mosaic” was defined as follows: subsequently developed embryos were partial cells with normal chromosomes and the others with abnormal chromosomes either with aneuploidy or without genomic disorders of deletions and/or duplications, i.e., 46XN, + mosaic (22) (33%) or 46 XN, dup (16) (p13.3p13.13) (5.7 Mb) (mos, 50%).

Because gametes’ chromosomal abnormalities should not be the source of “sole mosaic,” embryos with sole mosaic forms were considered mitotic chromosomal separation errors (Zhang X et al,.2021). Thus, in this study, “chromosomal-normal” and “sole mosaic” were included into the same group in the results.

“Sole aneuploidy” was defined as follows: subsequently developed embryos were aneuploid without any other errors, i.e., 47, XN, +22(×3).

“Sole deletion and/or duplication” indicates that embryos possess small (>10 Mb) or submicroscopic genomic deletions and/or duplications (1 kb to 10 Mb) without mosaic forms, e.g., 46, XN, dup (16) (p13.3p13.13) (5.7 Mb).

“Aneuploidy with errors” includes aneuploidy with any other solely chromosomal errors, i.e., 47, XN, +22(×3), dup (16) (p13.3p13.13) (5.7 Mb) or 47, XN, +22(×3), +mosaic (22) (33%).

“Euploidy with errors” includes “sole deletion and/or duplication” euploidy with any mosaic forms, i.e., 46, XN, dup (16) (p13.3p13.13) (5.7 Mb), +mosaic (22) (33%).

“Complex chromosomal errors” indicates aneuploidy with chromosomal deletion and/or duplication and mosaic forms, e.g., 45, XN, (−21), +4q (q12q31.1, ∼89 Mb, ×3), 9p (p20p21.1, ∼32 Mb,×1, mos, ∼50%).

The other classification of chromosomal errors was explored based on the coincidence of embryos’ and patients’ (the propositus) chromosomal/genomic abnormalities in PGT-SR. The results were grouped as “embryo’s chromosomal error coincident with female,” “embryo’s chromosomal error coincident with male,” “embryo’s chromosomal error inconsistent with female,” “embryo’s chromosomal error inconsistent with female,” “sole mosaic embryo,” and “chromosomal normal embryo.” Coincident errors mean the embryos’ chromosomes had complete or partial errors like female or male somatic chromosomes, i.e., 46, XX, t(1,16)(q42:q12) in somatic cells and 46, XN, +1q (q42.12→qter, ∼23.9 M, ×3), -16q (q12.1→q24.3, ∼39 M, ×1) in the embryo.

### Machine learning programming and statistical methods

PN machine learning, distinguishing female and male pronuclei, pronuclear labeling stability insurance, PN ranking order, and automatic mathematical model establishment were performed using Python 3.9.7 (downloaded from https://www.python.org/). Statistical analysis was performed using STATA12.0 software (Statacorp, TX, USA). Continuous variables are expressed as mean with standard deviation (SD) and categorical data are expressed as rate. The heterogeneity test for continuous variables, the Chi-square test for trend comparison, Spearman correlation analysis, multiple regression, and multiple logistic regression for relationships were used as appropriate.

## Results

After Mask R-CNN learning of 13419 handle-annotated 2PN images (276 embryos from 83 couples), the number of frames for AI automatic recognition reached 16634 for this sample. After comparison with handle annotation using 16634 AI automatic recognized images, 3343 images were more mislabeling than the actual 2PN images. In these 3343 images, 3192 images (95.48%) were came from early-stage PNs (12 hours post-insemination), the rest were came from 12-14 hours post-insemination and no images from 14 hours post-insemination. Additionally, 128 images were missing because of partial overlap of two PNs in middle- or late-stage PNs (14–22 hours post-insemination) (Figure 3B).

The accuracy of distinguish PN numbers by Mask R-CNN learning for (0, 1, 2, 3) reached 80.06% (13419/(13419+3342+128)) in recognition of all PN stages, 97.9% (13419/(13419+3342−3192+128)) in recognition at 12 hours post-insemination to PN disappearance, and 99.06% (13419/(13419+128)] at 14 hours to PN disappearance. No error was found in AI boundary drawing of PNs after handle checking except for PN number recognition-related boundary errors (e.g., mismarking vacuoles as PNs). For above errors in female and male pronucleus coefficient β1 calculation, original, adjusted (first and last three images deleted due to the high inaccuracy rate in recognition and high weight in the fitting curve model), 12 hours post-insemination, and 14 hours post-insemination values were extracted for effective testing. Then, 2146 embryos from 184 patients who have top-quality blastocysts for PGT were included in the data analysis. Different grades of PN identification are shown in Figure 3C. In total, 529 from 2146 zygotes (24.65%) underwent handle male/female PN reversal after computer marking and embryologist checking.

**Fig.3.**
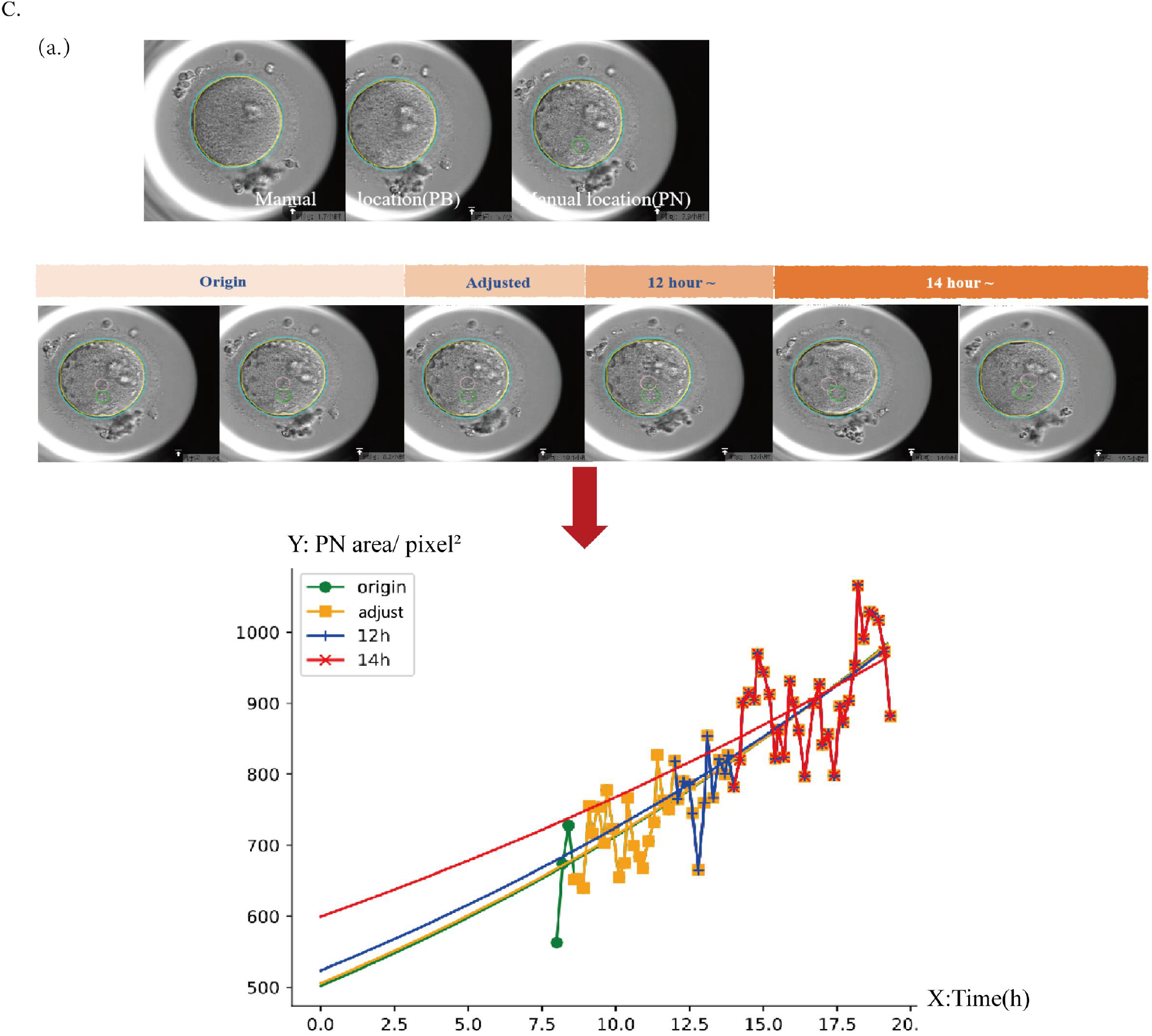
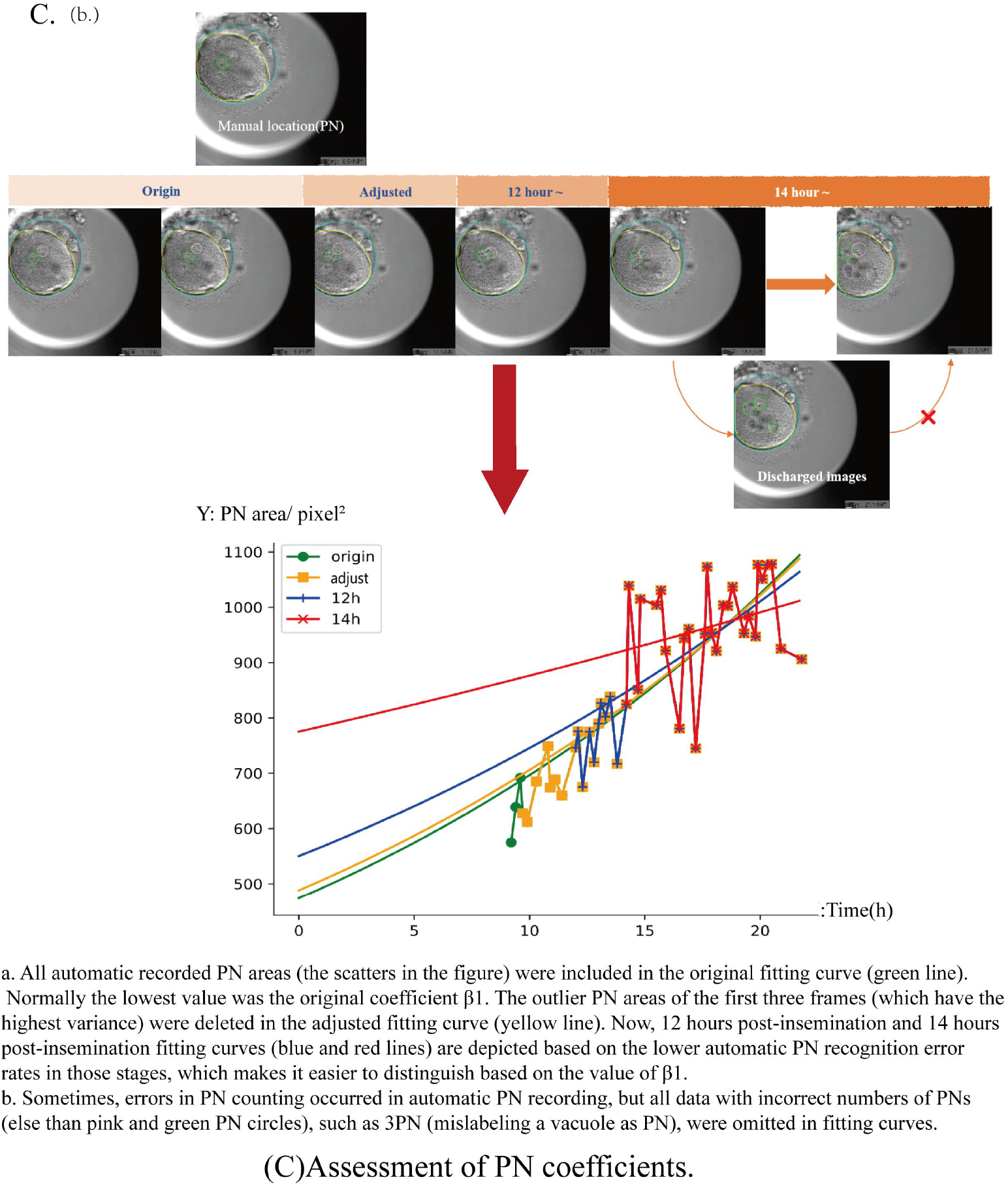
The statement of PN develpmental model and adjustment of coefficient

The baseline of patients and their IVF outcomes is shown in Table 1. Total frames of clear and distinguished 1655 2PN embryos reached 108587 images and these images were included to explore the factors that impacted female and male PN areas. In total, 1644 embryos with 108028 images were included for blastocyst formation analysis. Finally, 790 embryos with 52479 images from 155 patients were included for the analysis of both areas of pronuclei and PGT results.

**Table 1.** Baseline characteristics of patients.

| Patient characteristics | Mean (SD) |
| --- | --- |
| Age at IVF cycle | 30.55 ± 3.94 |
| Retrieval oocyte | 16.79 ± 6.81 |
| AMH level | 4.30 ± 2.77 |
| FSH (basic) | 5.59 ± 2.60 |
| BMI | 21.77 ± 2.76 |
| Gn day | 9.26 ± 1.56 |
| Gn dose | 1875.08 ± 622.27 |
| MII oocyte | 14.09 ± 6.20 |
| Infertility diagnosis (%) |  |
| Decreased ovarian reserve | 2.22% |
| Habitual loss | 9.63% |
| Male chromosome abnormality | 35.64% |
| Female chromosome abnormality | 31.92% |
| Uterine/fallopian tube factor | 8.00% |
| Obstetric abnormality | 5.31% |
| Unexplained | 7.28% |
| IVF protocol (%) |  |
| Gn-a | 43.1% |
| Gn-ant | 56.9% |

No clinical or cell biological factors were correlated with the female pronucleus coefficient β1 except for the male pronucleus coefficient β1 (r = 0.75, *P* < 0.01, Table 2) and the distribution data of the pronuclear area showed significant heterogeneity in individual patients. This heterogeneity makes it impossible to find a normal range of coefficient β1 (original data: Q=96.32, df=183, I^2^=95.6%, Supplement Figure 1). For homogenized exploration of the correlation between pronucleus area coefficient β1 and PGT results, ranking orders (biggest to smallest, Supplement Figure 2) of tmale/female were employed for testing in every patient. In blastocyst formation analysis, 1064 zygotes successfully developed into blastocysts and 580 failed; no significant relationship was observed between ranking order of the female or male pronucleus coefficient β1 and blastocyst formation (Table 3).

**Table 2.** The correlation between the male and female pronuclear coefficient β1.

| Dependent variable: female<br>pronuclear( coefficient $\beta 1$) | standard $\beta$ | *P* |
| --- | --- | --- |
| male pronuclear | 0.75 | 0.000 |
| embryo | -0.06 | 0.064 |
| Gn day | -0.08 | 0.092 |
| Gn dose | 0.08 | 0.051 |
| MII oocytes number | 0.04 | 0.232 |
| AMH | 0.02 | 0.409 |
| BMI | 0.07 | 0.200 |
| FSH (basic) | -0.01 | 0.755 |
| Female age | -0.04 | 0.095 |
| Infertility diagnosis | -0.02 | 0.322 |
| IVF protocol | 0.02 | 0.496 |

**Table 3.**
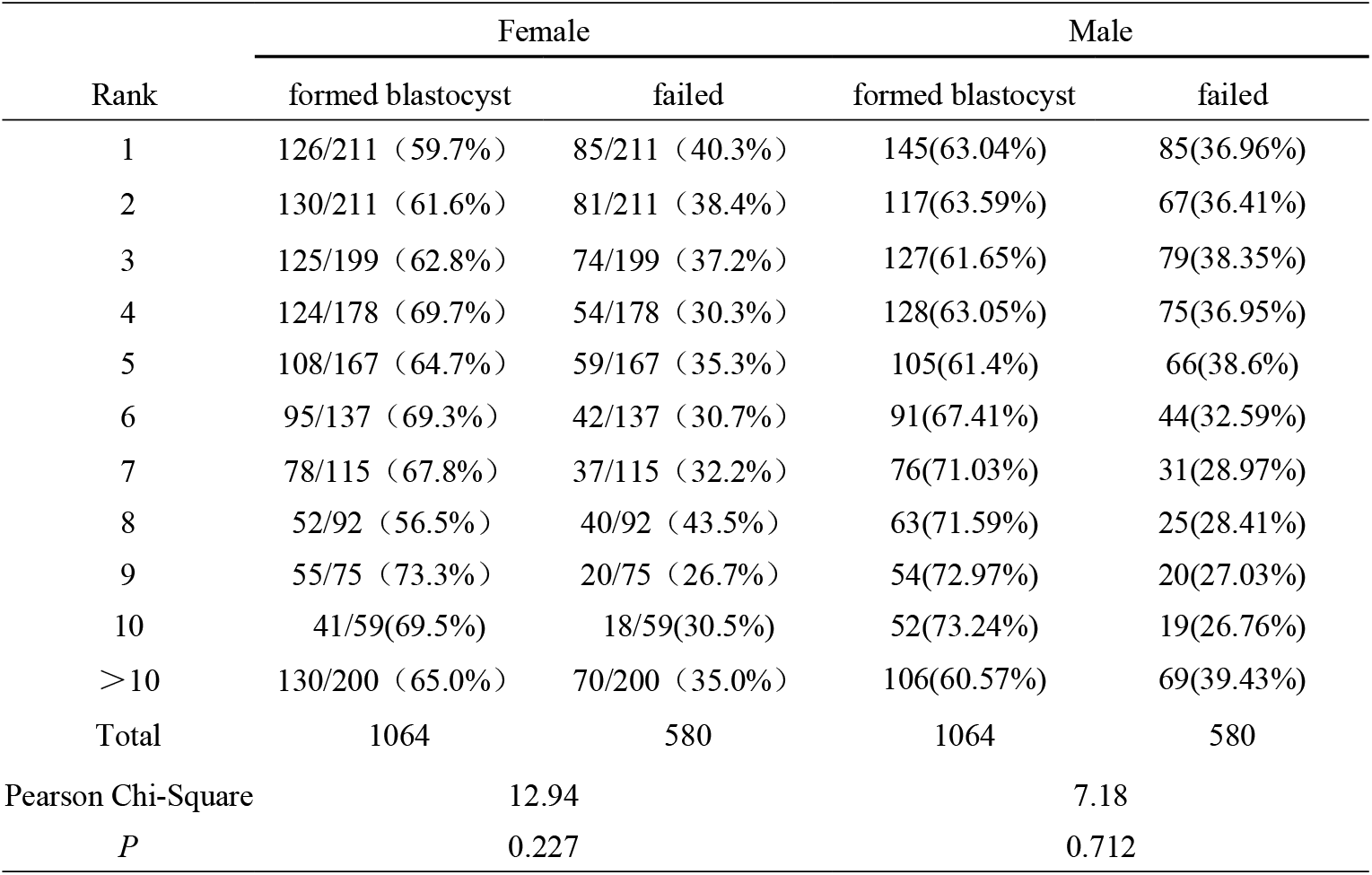
The chromosome-normal rate between blastocysts and embryos that failed to form blastocysts based on the rank of the pronucleus area coefficient β1.

| Rank | Female |  | Male |  |
| --- | --- | --- | --- | --- |
|  | formed blastocyst | failed | formed blastocyst | failed |
| 1 | 126/211 (59.7%) | 85/211 (40.3%) | 145(63.04%) | 85(36.96%) |
| 2 | 130/211 (61.6%) | 81/211 (38.4%) | 117(63.59%) | 67(36.41%) |
| 3 | 125/199 (62.8%) | 74/199 (37.2%) | 127(61.65%) | 79(38.35%) |
| 4 | 124/178 (69.7%) | 54/178 (30.3%) | 128(63.05%) | 75(36.95%) |
| 5 | 108/167 (64.7%) | 59/167 (35.3%) | 105(61.4%) | 66(38.6%) |
| 6 | 95/137 (69.3%) | 42/137 (30.7%) | 91(67.41%) | 44(32.59%) |
| 7 | 78/115 (67.8%) | 37/115 (32.2%) | 76(71.03%) | 31(28.97%) |
| 8 | 52/92 (56.5%) | 40/92 (43.5%) | 63(71.59%) | 25(28.41%) |
| 9 | 55/75 (73.3%) | 20/75 (26.7%) | 54(72.97%) | 20(27.03%) |
| 10 | 41/59(69.5%) | 18/59(30.5%) | 52(73.24%) | 19(26.76%) |
| >10 | 130/200 (65.0%) | 70/200 (35.0%) | 106(60.57%) | 69(39.43%) |
| Total | 1064 | 580 | 1064 | 580 |
| Pearson Chi-Square | 12.94 |  | 7.18 |  |
| *P* | 0.227 |  | 0.712 |  |

PGT results from transferable blastocysts have a U curve in the female age-dependent distribution (Supplement Figure 3). By the female PN original coefficient β1 ranking, chromosome normal rate (“chromosomal-normal” and “sole mosaic” proportion in total embryos) in the blastocyst with biggest PN area (top 1) is much higher than that of the blastocyst with smallest PN (last 1) (58.06% vs. 45.16%, OR = 1.68 [1.07–2.64]; *P* = 0.031), and the chromosome-normal rate of the top 2 blastocysts is higher than that of the last 2 blastocysts but with no statistical difference (59.31% vs. 50.49%; *P* = 0.091) (Supplemental Table 1-1). After adjusting for coefficient β1, the chromosome-normal rate in the top 1 blastocyst is much higher than that of the last blastocyst (58.71% vs. 45.16%, OR = 1.73 [1.10–2.71]; *P* = 0.023), and the chromosome-normal rate of the top 2 blastocysts is higher than that of the last 2 blastocysts but with no statistical difference (57.35% vs. 50.00%; *P* = 0.164) (Supplemental Table 1-2). For coefficient β1 12 hours post-insemination, the chromosome-normal rate in the top blastocyst is much higher than that of the last blastocyst (64.52% vs. 43.23%, OR = 2.39 [1.51–3.77]; *P* < 0.001], and the chromosome-normal rate in the top 2 blastocysts is higher than that of the last 2 blastocysts (63.73% vs. 47.55%, OR = 1.94 [1.30–2.88]; *P* = 0.001) (Supplemental Table 1-3). For coefficient β1 14 hours post-insemination, the chromosome-normal rate in the ranking top blastocyst is much higher than that of the last blastocyst (66.45% vs. 42.58%, OR = 2.61 [1.68–4.24]; *P* < 0.001), and the chromosome-normal rate in the top 2 blastocysts is higher than that of the last 2 blastocysts (64.22% vs. 48.04%, OR = 1.94 [1.31–2.89]; *P* = 0.001) (Supplemental Table 1-4). The trend that the top blastocysts showed higher chromosome-normal rates can be observed in Table 4 and Figure 4A. However, for the male PN coefficient β1, no significant difference was observed (Figure 4B).

**Table 4.** The rate of chromosome-normal blastocysts by the female PN coefficient β1.

| Rank | Origin | Adjusted | 12 h | 14 h |
| --- | --- | --- | --- | --- |
| 1 | 90/155 (58.06%) | 90/155 (58.06%) | 99/155 (63.87%) | 102/155 (65.81%) |
| 2 | 79/155 (50.97%) | 75/155 (48.38%) | 82/155 (52.9%) | 79/155 (50.97%) |
| 3 | 78/138 (56.52%) | 82/138 (59.42%) | 69/138 (50%) | 70/138 (50.72%) |
| 4 | 46/107 (42.99%) | 47/107 (43.93%) | 50/107 (46.73%) | 49/107 (45.79%) |
| 5 | 40/81 (49.38%) | 39/81 (48.15%) | 37/81 (45.68%) | 40/81 (49.38%) |
| 6 | 33/56 (58.93%) | 33/56 (58.93%) | 31/56 (55.36%) | 29/56 (51.79%) |
| 7 | 23/37 (62.16%) | 23/37 (62.16%) | 17/37 (45.95%) | 23/37 (62.16%) |
| 8 | 13/25 (52%) | 14/25 (56%) | 17/25 (68%) | 14/25 (56%) |
| >8 | 16/36 (44.44%) | 15/36 (41.67%) | 16/36 (44.44%) | 12/36 (33.33%) |

**Fig.4.**
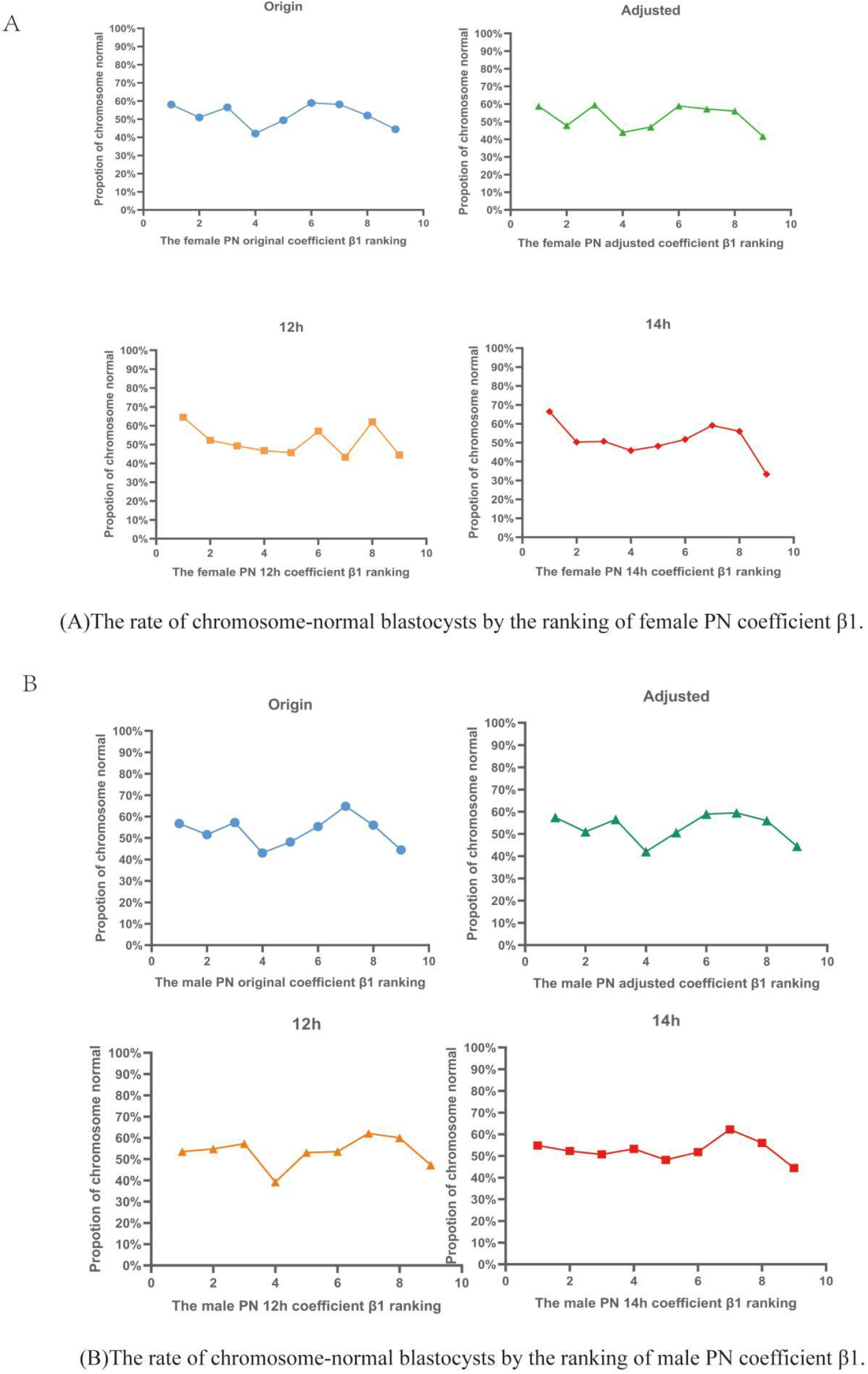
The rate of chromosome-normal blastocysts by the ranking of PN coefficient β1.

In all patients without distinguishing PGT-A and PGT-SR, the first and last ranking order could be used to detect the chromosome-normal and sole mosaic embryos (46.45% vs. 28.39% in chromosome-normal rate, 66.45% vs. 42.58% in chromosome-normal plus mosaic rate) (Figure 5A, Supplemental Table 2). No significant differences of chromosomal status were obtained by coefficient β1 ranking in the PGT-A population (Figure 5B, Supplemental Table 4-5). However, in the PGT-SR population, the first order embryos in coefficient β1 ranking have higher chromosome-normal and sole mosaic rates than the last embryos (40.54% vs. 21.62% in chromosome-normal rate, 66.22% vs. 32.43% in chromosome-normal plus mosaic rate) (Figure 5C, Supplemental Table 6). Relatively smaller chromosomal errors defined as “euploid with errors” and “sole deletion and/or duplication” were significantly different between first-order and last-order embryos, even with median-order embryos having a hierarchical difference (PGT 9.68% vs. 15.41% vs. 20%, PGT-SR 13.51% vs. 26.92% vs. 36.49%, respectively; *P* < 0.001) (Figure 5A and C, Supplemental Table 3, 6 and 7). For population- and chromosomal error-stratified analysis of coefficient β1 at 14 hours post-insemination, PGT-A and PGT-SR, “aneuploidy with errors” and “euploidy with errors” (and a more detailed error classification including “chromosomal-normal,” “sole mosaic,” “sole aneuploidy,” and mosaic forms, “sole deletion and/or duplication” and mosaic forms, and “complex chromosomal errors”), “embryo’s chromosomal error coincident/inconsistent with female” and “embryo’s chromosomal error coincident/inconsistent with male” are shown in Figures 5B and 5C (Supplemental Table 4-9). From the female propositus and embryo analysis in PGT-SR, coefficient β1 ranking has detection power in both coincident and inconsistent chromosomal errors (28.57% vs. 50%; 9.52% vs. 26.19%, *P*<0.05 respectively. Figure 5C, Supplemental Table 8), which implied inherited and novel errors in embryos, but no significant detection ability in male propositus and embryos (Figure 5C, Supplemental Table 9).

**Fig.5.**
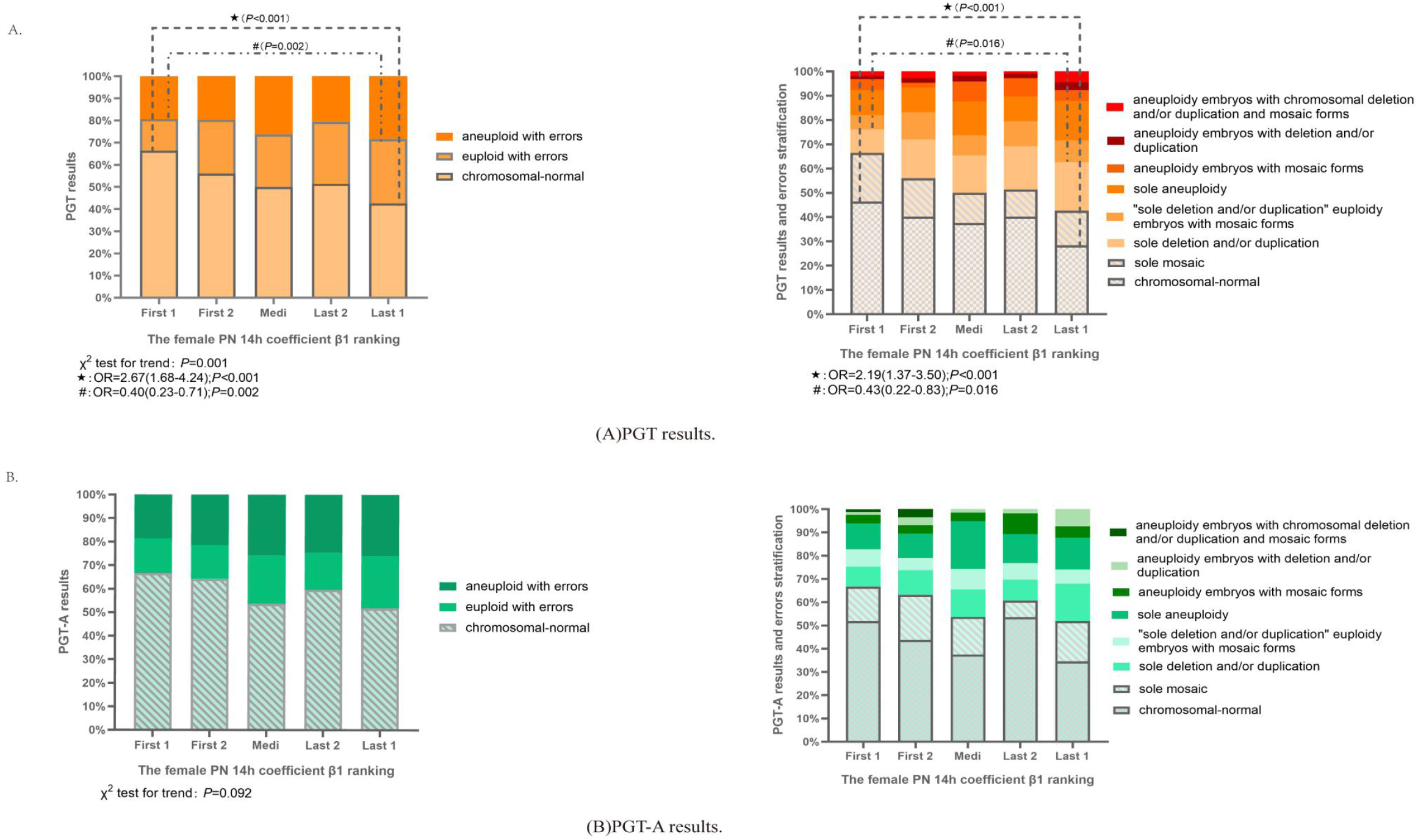
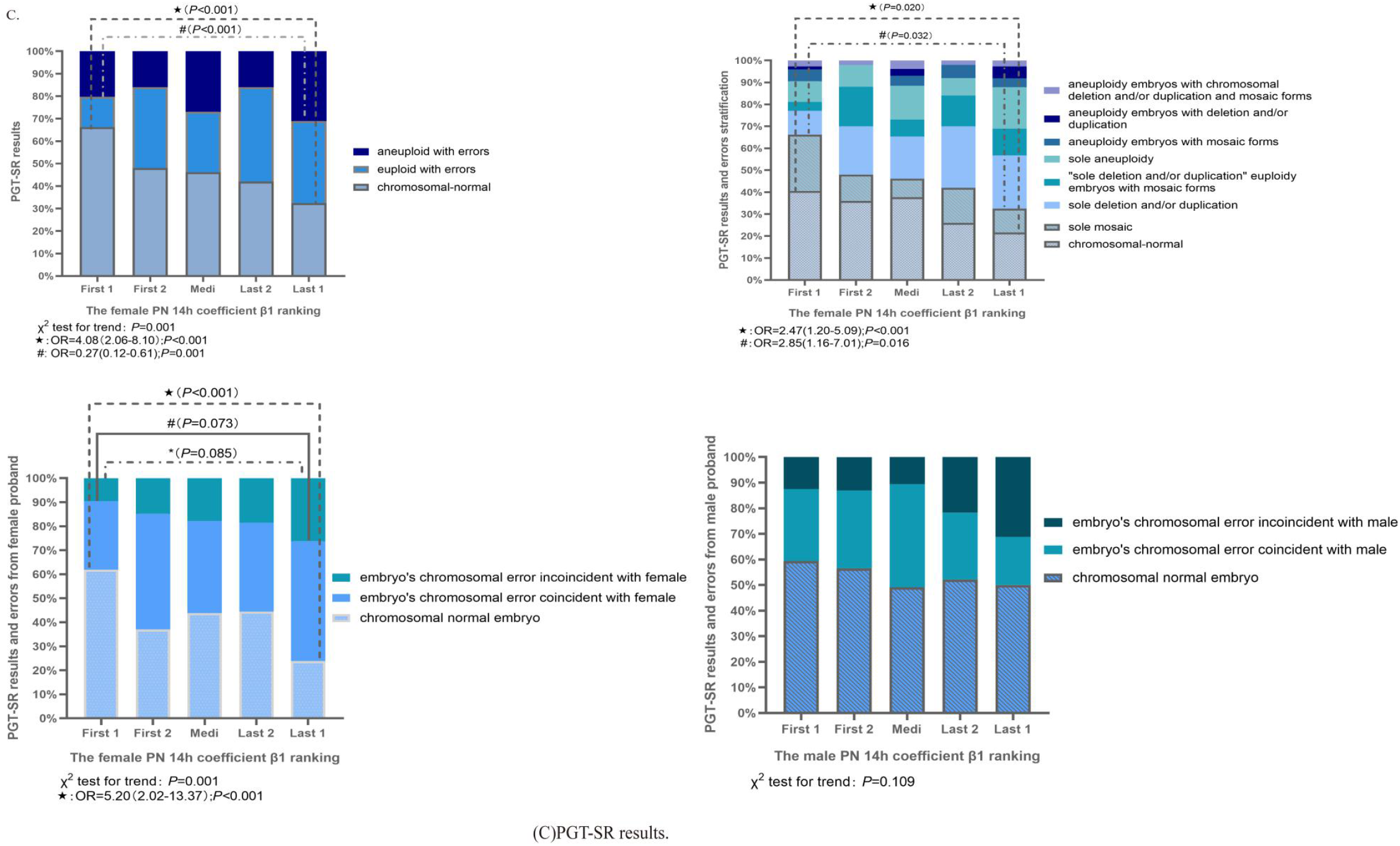
The correlation between PGT results and PNs coefficient β I ranking

## Discussion

An obvious relationship has been obtained between female PN and chromosome-normal rate in blastocyst-stage embryos for both original and adjusted analysis, but not for male PN. In the stratified analysis, female PN in the PGT-SR group, but no in the PGT-A group, have unambiguous detection power to distinguish relatively small chromosomal errors, such as “deletion and/or duplication” and mosaic forms. Inherited and novel errors in embryos could be found using female PN ranking in female diagnosis of the PGT-SR group. The overall positive pool effect of female PN diagnosis of chromosomal errors might be caused by the PGT-SR subgroup. The negative result in PGT-A might be because of a high false-positive rate (abnormal TE but normal ICM) as well as false-negative rate (normal TE but abnormal ICM) in this technique (Gleicher N et al., 2021).

From the PGT result, a high coincident U curve has been found as previously reported (Gruhn JR et al., 2019), but a small difference is that our age-distributed samples were blastocyst-stage embryos, not oocytes. Thus, chromosomal errors occurred post-PN from the cleavage to the morula and the blastocyst stage, and potential embryo self-correction in a later stage could reduce the power of PN predictors (Coticchio G et al., 2019;Grau N et al., 2011;Orvieto R et al., 2020).

The results indicated that embryonic chromosomal abnormality is more likely to be caused by eggs, especially in meiosis, and female PN developmental quantification could unveil the potential correlation (Miller MP et al., 2013;Warburton D et al., 1997;Mikwar M et al., 2020;Bolcun-Filas E et al., 2018;Webster A et al., 2017;Cairo G et al., 2020;Capalbo A et al., 2017). Again, the results confirmed the theory that the pronuclear stage could be the only road that mirrors the internal oocyte quality and chromosomal integrity. However, due to the low chromosomal error rate in sperm, male PNs had no predictive value (Bell AD et al., 2020). Here, we excluded chromosomal mosaicism because the typical mitotic errors could not be associated with PN stage, but the effects of mitotic errors will merge in cleavage- and blastocyst-stage embryos (Zhang X et al., 2021). In the earliest design of outcome measurements, the total chromosomal substance was classified as normal, deletion, and duplication, but no significant difference was obtained (Supplemental Table 10). Interestingly, when the outcome measure was changed into normal and abnormal, a clear difference was observed. The exact reason will be studied in further research.

A higher correlation has been obtained between female and male PN coefficient β1, but no clinical or cell biological factor exhibited a similar correlation in subsequent analyses. The high heterogeneity of the PN coefficient β1 made it impossible to establish a normal and abnormal range in clinical practice. No relationship between female or male PN and blastocyst formation has been found, revealing that protein and energy storage could be more important to the developmental viability of embryos than chromosomal normality, at least if chromosomal errors are not too big (Coticchio G et al., 2021).

This study was the first report on automatic calculation in morphologic quantitative data extraction by expert experience deep learning in human embryos. The results of the PN coefficient β1 suggest that detailed analysis of the images of developing embryos could improve our understanding of developmental biology, but more features of annotated embryos increase the errors in first and second polar body recognition (Cavazza T et al., 2021;Manor D et al., 1999; Otsuki J et al., 2017;Borges EJ et al., 2005). Further high-quality design studies are needed to improve the availability of quantitative PN assessment in clinical practice.

### Ideas and Speculation

Previously studies have reported dark box algorithm employed for embryo assessment compared with handle in this paper, but it could not explain how AI renders a decision from the embryos’ images. Embryo assessment from another access: embryo features deep learning and transfer those features into quantitative parameters for subsequent algorithm established and analyzed could more comprehensible for developmental biology and genetics. Then the PN morphology could mirror the internal quality of the chromosomal integrity of the oocyte and the spermatozoon.

## Reference

Alpha Scientists in Reproductive M, Embryology ESIGo. The Istanbul consensus workshop on embryo assessment: proceedings of an expert meeting. Human reproduction. 2011;26(6):1270–83.

Aydin S, Cinar O, Demir B, Korkmaz C, Ozdegirmenci O, Dilbaz S, et al. Is pronuclear scoring a really good predictor for ICSI cycles? Gynecological endocrinology : the official journal of the International Society of Gynecological Endocrinology. 2011;27(10):742–7.

Balaban B, Urman B, Isiklar A, Alatas C, Aksoy S, Mercan R, et al. The effect of pronuclear morphology on embryo quality parameters and blastocyst transfer outcome. Human reproduction (Oxford, England). 2001;16(11):2357–61.

Bar-Yoseph H, Levy A, Sonin Y, Alboteanu S, Levitas E, Lunenfeld E, et al. Morphological embryo assessment: reevaluation. Fertility and sterility. 2011;95(5):1624–8 e1-2.

Belkin, M., and P. Niyogi. “Towards a Theoretical Foundation for Laplacian-Based Manifold Methods.” International Conference on Computational Learning Theory Springer, Berlin, Heidelberg, 2005.

Bell AD, Mello CJ, Nemesh J, Brumbaugh SA, Wysoker A, McCarroll SA. Insights into variation in meiosis from 31,228 human sperm genomes. Nature. 2020;583(7815):259–64.

Bolcun-Filas E, Handel MA. Meiosis: the chromosomal foundation of reproduction. Biology of reproduction. 2018;99(1):112–26.

Borges EJ, Rossi LM, Farah L, Guilherme P, Rocha CC, Ortiz V, Iaconelli AJ. The impact of pronuclear orientation to select chromosomally normal embryos. J Assist Reprod Genet. 2005; 22(3):107–14. doi: 10.1007/s10815-005-4874-x. PMID: 16018240; PMCID: PMC3455178.

Cai D, He X, Han J, Zhang H-J. Orthogonal laplacianfaces for face recognition. IEEE Trans Image Process. 2006;15(11):3608–14.

Cairo G, Lacefield S. Establishing correct kinetochore-microtubule attachments in mitosis and meiosis. Essays Biochem. 2020 Sep 4;64(2):277–287.

Capalbo A, Hoffmann ER, Cimadomo D, Ubaldi FM, Rienzi L. Human female meiosis revised: new insights into the mechanisms of chromosome segregation and aneuploidies from advanced genomics and time-lapse imaging. Human reproduction update. 2017;23(6):706–22.

Cavazza T, Takeda Y, Politi AZ, Aushev M, Aldag P, Baker C, Choudhary M,Bucevičius J, Lukinavičius G, Elder K, Blayney M, Lucas-Hahn A, Niemann H,Herbert M, Schuh M. Parental genome unification is highly error-prone in mammalian embryos. Cell. 2021; 184(11):2860–2877.e22. doi:10.1016/j.cell.2021.04.013.

Chavez SL, Loewke KE, Han J, Moussavi F, Colls P, Munne S, et al. Dynamic blastomere behaviour reflects human embryo ploidy by the four-cell stage. Nature communications. 2012;3:1251.

Coticchio G, Fiorentino G, Nicora G, Sciajno R, Cavalera F, Bellazzi R, et al. Cytoplasmic movements of the early human embryo: imaging and artificial intelligence to predict blastocyst development. Reproductive biomedicine online. 2021;42(3):521–8.

Coticchio G, Lagalla C, Sturmey R, Pennetta F, Borini A. The enigmatic morula: mechanisms of development, cell fate determination, self-correction and implications for ART. Human reproduction update. 2019;25(4):422–38.

Coticchio G, Mignini Renzini M, Novara PV, Lain M, De Ponti E, Turchi D, et al. Focused time-lapse analysis reveals novel aspects of human fertilization and suggests new parameters of embryo viability. Human reproduction. 2018;33(1):23–31.

Daughtry BL, Rosenkrantz JL, Lazar NH, Fei SS, Redmayne N, Torkenczy KA, et al. Single-cell sequencing of primate preimplantation embryos reveals chromosome elimination via cellular fragmentation and blastomere exclusion. Genome Res. 2019;29(3):367–82.

Doody KJ. The time has come to reevaluate the fertilization check. Fertility and sterility. 2021;115(1):74–5.

Gámiz P, Rubio C, de los Santos MJ, Mercader A, Simón C, Remohí J, et al. The effect of pronuclear morphology on early development and chromosomal abnormalities in cleavage-stage embryos. Human reproduction (Oxford, England). 2003;18(11):2413–9.

Gianaroli L, Magli MC, Ferraretti AP, Lappi M, Borghi E, Ermini B. Oocyte euploidy, pronuclear zygote morphology and embryo chromosomal complement. Human reproduction. 2007;22(1):241–9.

Gleicher N, Patrizio P, Brivanlou A. Preimplantation Genetic Testing for Aneuploidy - a Castle Built on Sand. Trends in molecular medicine. 2021;27(8):731–42.

Grau N, Escrich L, Martin J, Rubio C, Pellicer A, Escriba MJ. Self-correction in tripronucleated human embryos. Fertility and sterility. 2011;96(4):951–6.

Gruhn JR, Zielinska AP, Shukla V, Blanshard R, Capalbo A, Cimadomo D, et al. Chromosome errors in human eggs shape natural fertility over reproductive life span. Science. 2019;365(6460):1466–9.

He K, Gkioxari G, Dollar P, Girshick R. Mask R-CNN. IEEE Trans Pattern Anal Mach Intell. 2020 Feb;42(2):386–397. doi: 0.1109/TPAMI.2018.2844175.

Hoffman R, Gross L. Modulation contrast microscope. Appl Opt. 1975;14(5):1169–76.

Jaroudi K, Al-Hassan S, Sieck U, Al-Sufyan H, Al-Kabra M, Coskun S. Zygote transfer on day 1 versus cleavage stage embryo transfer on day 3: a prospective randomized trial. Human reproduction (Oxford, England). 2004;19(3):645–8.

Kuliev A, Zlatopolsky Z, Kirillova I, Spivakova J, Cieslak Janzen J. Meiosis errors in over 20,000 oocytes studied in the practice of preimplantation aneuploidy testing. Reproductive biomedicine online. 2011;22(1):2–8.

Lamb NE, Freeman SB, Savage-Austin A, Pettay D, Taft L, Hersey J, et al. Susceptible chiasmate configurations of chromosome 21 predispose to non-disjunction in both maternal meiosis I and meiosis II. Nat Genet. 1996;14(4):400–5.

Li M, Huang J, Zhuang X, Lin S, Dang Y, Wang Y, et al. Obstetric and neonatal outcomes after the transfer of vitrified-warmed blastocysts developing from nonpronuclear and monopronuclear zygotes: a retrospective cohort study. Fertility and sterility. 2021;115(1):110–7.

Manor D, Drugan A, Stein D, Pillar M, Itskovitz-Eldor J. Unequal pronuclear size--a powerful predictor of embryonic chromosome anomalies. J Assist Reprod Genet. 1999;16(7):385–9. doi: 10.1023/a:1020550115345. PMID: 10459523;PMCID: PMC3455778.

Mikwar M, MacFarlane AJ, Marchetti F. Mechanisms of oocyte aneuploidy associated with advanced maternal age. Mutation research Reviews in mutation research. 2020;785:108320.

Miller MP, Amon A, Unal E. Meiosis I: when chromosomes undergo extreme makeover. Current opinion in cell biology. 2013;25(6):687–96.

Munné S. Chromosome abnormalities and their relationship to morphology and development of human embryos. Reproductive biomedicine online. 2006;12(2):234–53.

Nicoli A, Palomba S, Capodanno F, Fini M, Falbo A, La Sala GB. Pronuclear morphology evaluation for fresh in vitro fertilization (IVF) and intracytoplasmic sperm injection (ICSI) cycles: a systematic review. J Ovarian Res. 2013;6(1):64.

Orvieto R, Shimon C, Rienstein S, Jonish-Grossman A, Shani H, Aizer A. Do human embryos have the ability of self-correction? Reproductive biology and endocrinology : RB&E. 2020;18(1):98.

Otsuki J, Iwasaki T, Tsuji Y, Katada Y, Sato H, Tsutsumi Y, Hatano K, Furuhashi K, Matsumoto Y, Kokeguchi S, Shiotani M. Potential of zygotes to produce live births can be identified by the size of the male and female pronuclei just before their membranes break down. Reprod Med Biol. 2017 ;16(2):200–205. doi: 10.1002/rmb2.12032. PMID: 29259470; PMCID: PMC5661814

Rienzi L, Ubaldi F, Iacobelli M, Ferrero S, Minasi MG, Martinez F, et al. Day 3 embryo transfer with combined evaluation at the pronuclear and cleavage stages compares favourably with day 5 blastocyst transfer. Human reproduction (Oxford, England). 2002;17(7):1852–5.

Roos Kulmann MI, Lumertz Martello C, Bos-Mikich A, Frantz N. Pronuclear and blastocyst morphology are associated age-dependently with embryo ploidy in in vitro fertilization cycles. Human fertility. 2020:1–8.

Scheffler K, Uraji J, Jentoft I, Cavazza T, Monnich E, Mogessie B, et al. Two mechanisms drive pronuclear migration in mouse zygotes. Nature communications. 2021;12(1):841.

Scott LA, Smith S. The successful use of pronuclear embryo transfers the day following oocyte retrieval. Human reproduction (Oxford, England). 1998;13(4):1003–13.

Tesarik J, Junca AM, Hazout A, Aubriot FX, Nathan C, Cohen-Bacrie P, et al. Embryos with high implantation potential after intracytoplasmic sperm injection can be recognized by a simple, non-invasive examination of pronuclear morphology. Human reproduction (Oxford, England). 2000;15(6):1396–9.

Warburton D. Human female meiosis: new insights into an error-prone process. Am J Hum Genet. 1997;61(1):1–4.

Webster A, Schuh M. Mechanisms of Aneuploidy in Human Eggs. Trends in cell biology. 2017;27(1):55–68.

Wiker S, Malter H, Wright G, Cohen J. Recognition of paternal pronuclei in human zygotes. J In Vitro Fert Embryo Transf. 1990;7(1):33–7.

Wright G, Wiker S, Elsner C, Kort H, Massey J, Mitchell D, et al. Observations on the morphology of pronuclei and nucleoli in human zygotes and implications for cryopreservation. Human reproduction (Oxford, England). 1990;5(1):109–15.

Yang D, Feng D, Gao Y, Sagnelli M, Wang X, Li D. An effective method for trophectoderm biopsy using mechanical blunt dissection: a step-by-step demonstration. Fertility and sterility. 2020;114(2):438–9.

Zhang X, Yang J, Han W, Li C, Huang G. Blastomere movement correlates with ploidy and mosaicism in early-stage human embryos after in vitro fertilization. Zygote. 2021:1–15.

Zhao M, Xu M, Li H, Alqawasmeh O, Chung JPW, Li TC, et al. Application of convolutional neural network on early human embryo segmentation during in vitro fertilization. Journal of cellular and molecular medicine. 2021;25(5):2633–44.

Zollner U, Zollner KP, Hartl G, Dietl J, Steck T. The use of a detailed zygote score after IVF/ICSI to obtain good quality blastocysts: the German experience. Human reproduction (Oxford, England). 2002;17(5):1327–33.

