## Supplemental Tables and Figures for "Variation of female pronucleus reveal oocyte or embryo abnormality: An expert experience deep learning of non-dark box analysis"

Supplement

Table 1-1. Finding a chromosome-normal blastocyst by the original ranking of embryos.

| Number of embryo | N | First | Median 1 | Last | First 1 & 2 | Median 2 | Last 1 & 2 |
|---|---|---|---|---|---|---|---|
| 2 | 34 | 11/17 (64.71) | / | 7/17 (41.18) | / | / | / |
| 3 | 93 | 16/31 (51.61) | 14/31 (45.16) | 18/31 (58.06) | / | / | / |
| 4 | 104 | 11/26 (42.31) | 26/52 (50.00) | 10/26 (38.46) | 21/52 (40.38) | / | 26/52 (50.00) |
| 5 | 125 | 12/25 (48.00) | 36/75 (48.00) | 10/25 (40.00) | 25/50 (50.00) | 13/25 (52.00) | 20/50 (40.00) |
| 6 | 114 | 15/19 (78.95) | 42/76 (55.26) | 7/19 (36.84) | 28/38 (73.68) | 19/38 (50.00) | 17/38 (44.74) |
| 7 | 84 | 11/12 (91.67) | 39/60 (65.00) | 8/12 (66.67) | 21/24 (87.50) | 20/36 (55.56) | 17/24 (70.83) |
| 8 | 96 | 8/12 (66.67) | 40/72 (55.56) | 6/12 (50.00) | 14/24 (58.33) | 27/48 (56.25) | 13/24 (54.17) |
| 9 | 9 | 0/1 (0) | 3/7 (42.86) | 0/1 (0) | 1/2 (50.00) | 2/5 (40.00) | 0/2 (0) |
| 10 | 60 | 3/6 (50.00) | 32/48 (66.67) | 3/6 (50.00) | 6/12 (50.00) | 25/36 (69.44) | 7/12 (58.33) |
| 11 | 22 | 1/2 (50.00) | 8/18 (33.33) | 1/2 (50.00) | 2/4 (50.00) | 6/14 (42.86) | 2/4 (50.00) |
| 12 | 36 | 2/3 (66.67) | 10/30 (33.33) | 0/3 (0) | 3/6 (50.00) | 8/24 (33.33) | 1/6 (16.67) |
| 13 | 13 | 0/1 (0.00) | 7/11 (63.64) | 0/1 (0) | 0/2 (0.00) | 7/9 (77.78) | 0/2 (0) |
| Total | 790 | 90/155 (58.06)* | 227/480 (47.29) | 70/155 (45.16)* | 121/204 (59.31)# | 127/235 (54.04) | 103/204 (50.49)# |

*58.06% vs. 45.16%, OR = 1.68 (1.07–2.64); *P* = 0.031

#59.31% vs. 50.49%, OR = 1.43 (0.97–2.14); *P* = 0.091

Table 1-2. Finding a chromosome-normal blastocyst by the adjusted ranking of embryos.

| Number of embryo | N | First | Median 1 | Last | First 2 | Median 2 | Last 2 |
|---|---|---|---|---|---|---|---|
| 2 | 34 | 10/17 (58.82) | | 8/17 (47.06) | / | / | / |
| 3 | 93 | 17/31 (54.84) | 13/31 (41.94) | 18/31 (58.06) | / | / | / |
| 4 | 104 | 12/26 (46.15) | 27/52 (51.92) | 8/26 (30.77) | 21/52 (40.38) | / | 26/52 (50.00) |
| 5 | 125 | 13/25 (52.00) | 35/75 (46.67) | 10/25 (40.00) | 25/50 (50.00) | 14/25 (56.00) | 19/50 (38.00) |
| 6 | 114 | 14/19 (73.68) | 42/76 (55.26) | 8/19 (42.11) | 26/38 (68.42) | 21/38 (55.26) | 17/38 (44.74) |
| 7 | 84 | 11/12 (91.67) | 39/60 (65.00) | 8/12 (66.67) | 20/24 (83.33) | 21/36 (58.33) | 17/24 (70.83) |
| 8 | 96 | 8/12 (66.67) | 40/72 (55.56) | 6/12 (50.00) | 15/24 (62.50) | 26/48 (54.17) | 13/24 (54.17) |
| 9 | 9 | 0/1 (0) | 3/7 (42.86) | 0/1 (0) | 0/2 (0) | 3/5 (60.00) | 0/2 (0) |
| 10 | 60 | 3/6 (50.00) | 32/48 (66.67) | 3/6 (50.00) | 6/12 (50.00) | 25/36 (69.44) | 7/12 (58.33) |
| 11 | 22 | 1/2 (50.00) | 8/18 (44.44) | 1/2 (50.00) | 1/4 (25.00) | 7/14 (50.00) | 2/4 (50.00) |
| 12 | 36 | 2/3 (66.67) | 10/30 (33.33) | 0/3 (0) | 3/6 (50.00) | 8/24 (33.33) | 1/6 (16.67) |
| 13 | 13 | 0/1 (0) | 7/11 (63.64) | 0/1 (0) | 0/2 (0) | 7/9 (77.78) | 0/2 (0) |
| Total | 790 | 91/155 (58.71)* | 246/480 (51.25) | 70/155 (45.16)* | 117/204 (57.35)# | 132/235 (56.17) | 102/204 (50.00)# |

*58.71% vs. 45.16%, OR = 1.73 (1.10–2.71); *P* = 0.023

#57.35% vs. 50.00%, OR = 1.35 (0.91–1.99); *P* = 0.164

Table 1-3. Finding a chromosome-normal blastocyst by the ranking of 12 h embryos.

| Number of embryo | N | First | Median 1 | Last | First 2 | Median 2 | Last 2 |
|---|---|---|---|---|---|---|---|
| 2 | 34 | 11/17 (64.71) | | 7/17 (41.18) | / | / | / |
| 3 | 93 | 19/31 (61.29) | 14/31 (45.16) | 15/31 (48.39) | / | / | / |
| 4 | 104 | 13/26 (50.00) | 26/52 (50.00) | 8/26 (30.77) | 26/52 (50.00) | / | 21/52 (40.38) |
| 5 | 125 | 15/25 (60.00) | 34/75 (45.33) | 9/25 (36.00) | 27/50 (54.00) | 12/25 (48.00) | 19/50 (38.00) |
| 6 | 114 | 14/19 (73.68) | 42/76 (55.26) | 8/19 (42.11) | 25/38 (65.79) | 20/38 (52.63) | 19/38 (50.00) |
| 7 | 84 | 12/12 (100.00) | 39/60 (65.00) | 7/12 (58.33) | 22/24 (91.67) | 20/36 (55.56) | 16/24 (66.67) |
| 8 | 96 | 9/12 (75.00) | 35/72 (48.61) | 10/12 (83.33) | 14/24 (58.33) | 27/48 (56.25) | 13/24 (54.17) |
| 9 | 9 | 1/1 (100.00) | 2/7 (28.57) | 0/1 (0) | 1/2 (50.00) | 2/5 (40.00) | 0/2 (0) |
| 10 | 60 | 4/6 (66.67) | 32/48 (66.67) | 2/6 (33.33) | 10/12 (83.33) | 22/36 (61.11) | 6/12 (50.00) |
| 11 | 22 | 0/2 (0) | 9/18 (50.00) | 1/2 (50.00) | 2/4 (50.00) | 5/14 (35.71) | 3/4 (75.00) |
| 12 | 36 | 2/3 (66.67) | 10/30 (33.33) | 0/3 (0) | 3/6 (50.00) | 9/24 (37.50) | 0/6 (0) |
| 13 | 13 | 0/1 (0) | 7/11 (63.64) | 0/1 (0) | 0/2 (0) | 7/9 (77.78) | 0/2 (0) |
| Total | 790 | 100/155 (64.52)* | 250/480 (52.08) | 67/155 (43.23)* | 130/204 (63.73)# | 124/235 (52.77) | 97/204 (47.55)# |

*64.52% vs. 43.23%, OR = 2.39 (1.51–3.77); $P < 0.001$

#63.73% vs. 47.55%, OR = 1.94 (1.30–2.88); $P = 0.001$

Table 1-4. Finding a chromosome-normal blastocyst by the ranking of 14 h embryos.

| Number of embryo | N | First | Median 1 | Last | First 2 | Median 2 | Last 2 |
|---|---|---|---|---|---|---|---|
| 2 | 34 | 11/17 (64.71) | | 7/17 (41.18) | / | / | / |
| 3 | 93 | 21/31 (67.74) | 11/31 (35.48) | 16/31 (51.61) | / | / | / |
| 4 | 104 | 13/26 (50.00) | 25/52 (48.08) | 9/26 (34.62) | 26/52 (50.00) | / | 21/52 (40.38) |
| 5 | 125 | 13/25 (52.00) | 36/75 (48.00) | 9/25 (36.00) | 26/50 (52.00) | 11/25 (44.00) | 21/50 (42.00) |
| 6 | 114 | 17/19 (89.47) | 39/76 (51.32) | 8/19 (42.11) | 28/38 (73.68) | 17/38 (44.74) | 19/38 (50.00) |
| 7 | 84 | 11/12 (91.67) | 40/60 (66.67) | 7/12 (58.33) | 21/24 (87.50) | 23/36 (63.89) | 14/24 (58.33) |
| 8 | 96 | 9/12 (75.00) | 38/72 (52.78) | 7/12 (58.33) | 14/24 (58.33) | 26/48 (54.17) | 14/24 (58.33) |
| 9 | 9 | 1/1 (100.00) | 2/7 (28.57) | 0/1 (0) | 1/2 (50.00) | 2/5 (40.00) | 0/2 (0) |
| 10 | 60 | 5/6 (83.33) | 31/48 (64.58) | 2/6 (33.33) | 9/12 (75.00) | 24/36 (66.67) | 5/12 (41.67) |
| 11 | 22 | 0/2 (0) | 9/18 (50.00) | 1/2 (50.00) | 2/4 (50.00) | 6/14 (42.86) | 2/4 (50.00) |
| 12 | 36 | 2/3 (66.67) | 10/30 (33.33) | 0/3 (0) | 3/6 (50.00) | 8/24 (33.33) | 1/6 (16.67) |
| 13 | 13 | 0/1 (0) | 7/11 (63.64) | 0/1 (0) | 1/2 (50.00) | 5/9 (55.56) | 1/2 (50.00) |
| Total | 790 | 103/155 (66.45)* | 248/480 (51.67) | 66/155 (42.58)* | 131/204 (64.22)# | 122/235 (51.91) | 98/204 (48.04)# |

*66.45% vs. 42.58%, OR = 2.61 (1.68–4.24); $P < 0.001$

#64.22% vs. 48.04%, OR = 1.94 (1.31–2.89); $P = 0.001$

Table 2. PGT results and female PN 14 h coefficient β1 ranking.

| PGT result | First 1 | First 2 | Median | Last 2 | Last 1 |
|---|---|---|---|---|---|
| Chromosomal-normal and mosaic | 103/155 (66.45%)* | 60/107 (56.07%) | 133/266 (50%) | 55/107 (51.4%) | 66/155 (42.58%)* |
| Euploid with errors | 22/155 (14.19%)# | 26/107 (24.3%) | 63/266 (23.68%) | 30/107 (28.04%) | 45/155 (29.03%)# |
| Aneuploid with errors | 30/155 (19.35%) | 21/107 (19.63%) | 70/266 (26.32%) | 22/107 (20.56%) | 44/155 (28.39%) |

*66.45% vs. 42.58%, OR = 2.67 (1.68 – 4.24), $P < 0.001$;

#14.19% vs. 29.03%, OR = 0.40 (0.23 – 0.71), $P = 0.002$

Table 3. PGT results (errors stratification) and female PN 14 h coefficient β1 ranking.

| PGT result | First 1 | First 2 | Median | Last 2 | Last 1 |
|---|---|---|---|---|---|
| Chromosome-normal | 72/155 (46.45%)* | 43/107 (40.19%) | 100/266 (37.59%) | 43/107 (40.19%) | 44/155 (28.39%)* |
| Sole mosaic | 31/155 (20.00%) | 17/107 (15.89%) | 33/266 (12.41%) | 12/107 (11.21%) | 22/155 (14.19%) |
| Sole deletion and/or duplication | 15/155 (9.68%)# | 17/107 (15.89%) | 41/266 (15.41%) | 19/107 (17.76%) | 31/155 (20.00%)# |
| Sole deletion and/or duplication euploidy embryos with mosaic forms | 9/155 (5.81%) | 12/107 (11.21%) | 22/266 (8.27%) | 11/107 (10.28%) | 14/155 (9.03%) |
| Sole aneuploidy | 16/155 (10.32%) | 11/107 (10.28%) | 37/266 (13.91%) | 11/107 (10.28%) | 25/155 (16.13%) |
| Aneuploidy embryos with mosaic forms | 7/155 (4.52%) | 2/107 (1.87%) | 22/266 (8.27%) | 8/107 (7.48%) | 7/155 (4.52%) |
| Aneuploidy embryos with deletion and/or duplication | 2/155 (1.29%) | 2/107 (1.87%) | 6/266 (2.26%) | 2/107 (1.87%) | 5/155 (3.23%) |
| Aneuploidy embryos with chromosomal deletion and/or duplication and mosaic forms | 3/155 (1.94%) | 3/107 (2.8%) | 5/266 (1.88%) | 1/107 (0.93%) | 7/155 (4.52%) |

*46.45% vs. 28.39%, OR = 2.19 (1.37 – 3.50), $P < 0.001$;

#9.68% vs. 20%, OR = 0.43 (0.22–0.83), $P = 0.016$

Table 4. PGT-A results and female PN 14 h coefficient β1 ranking.

| PGT-A result | First 1 | First 2 | Median | Last 2 | Last 1 |
|---|---|---|---|---|---|
| Chromosomal-normal and mosaic | 54/81 (66.67%) | 36/56 (64.29%) | 73/136 (53.68%) | 34/57 (59.65%) | 42/81 (51.85%) |
| Euploid with errors | 12/81 (14.81%) | 8/56 (14.29%) | 28/136 (20.59%) | 9/57 (15.79%) | 18/81 (22.22%) |
| Aneuploid with errors | 15/81 (18.52%) | 12/56 (21.43%) | 35/136 (25.74%) | 14/57 (24.56%) | 21/81 (25.93%) |

Table 5. PGT-A results (errors stratification) and female PN 14 h coefficient β1 ranking.

| PGT-A result | First 1 | First 2 | Median | Last 2 | Last 1 |
|---|---|---|---|---|---|
| Chromosomal-normal | 42/81 (51.85%) | 25/57 (43.86%) | 51/136 (37.5%) | 30/57 (52.63%) | 28/81 (34.57%) |
| Sole mosaic | 12/81 (14.81%) | 11/57 (19.3%) | 22/136 (16.18%) | 4/57 (7.02%) | 14/81 (17.28%) |
| Sole deletion and/or duplication | 7/81 (8.64%) | 6/57 (10.53%) | 16/136 (11.76%) | 5/57 (8.77%) | 13/81 (16.05%) |
| Sole deletion and/or duplication euploidy embryos with mosaic | 6/81 (7.41%) | 3/57 (5.26%) | 12/136 (8.82%) | 4/57 (7.02%) | 5/81 (6.17%) |

| | | | | | |
|---|---|---|---|---|---|
| forms | | | | | |
| Sole aneuploidy | 9/81 (11.11%) | 6/57 (10.53%) | 28/136 (20.59%) | 7/57 (12.28%) | 11/81 (13.58%) |
| Aneuploidy embryos with mosaic forms | 3/81 (3.7%) | 2/57 (3.51%) | 5/136 (3.68%) | 5/57 (8.77%) | 4/81 (4.94%) |
| Aneuploidy embryos with deletion and/or duplication | 1/81 (1.23%) | 2/57 (3.51%) | 2/136 (1.47%) | 1/57 (1.75%) | 6/81 (7.41%) |
| Aneuploidy embryos with chromosomal deletion and/or duplication and mosaic forms | 1/81 (1.23%) | 2/57 (3.51%) | 0 | 0 | 0 |

Table 6. PGT-SR results and female PN 14 h coefficient β1 ranking.

| PGT-SR result | First 1 | First 2 | Median | Last 2 | Last 1 |
|---|---|---|---|---|---|
| Chromosome-normal and mosaic | 49/74 (66.22%)* | 24/50 (48%) | 60/130 (46.15%) | 21/50 (42%) | 24/74 (32.43%)* |
| Euploid with errors | 10/74 (13.51%)# | 18/50 (36%) | 35/130 (26.92%) | 21/50 (42%) | 27/74 (36.49%)# |
| Aneuploid with errors | 15/74 (20.27%) | 8/50 (16%) | 35/130 (26.92%) | 8/50 (16%) | 23/74 (31.08%) |

*66.22% vs. 32.43%, OR = 4.08 (2.06–8.10), $P < 0.001$

#13.51% vs. 36.49%, OR = 0.27 (0.12–0.61), $P = 0.001$

Table 7. PGT-SR results (errors stratification) and female PN 14 h coefficient β1 ranking.

| PGT-SR result | First 1 | First 2 | Median | Last 2 | Last 1 |
|---|---|---|---|---|---|
| Chromosome-normal | 30/74 (40.54%)* | 18/50 (36.00%) | 49/130 (37.69%) | 13/50 (26.00%) | 16/74 (21.62%)* |
| Sole mosaic | 19/74 (25.68%)# | 6/50 (12.00%) | 11/130 (8.46%) | 8/50 (16.00%) | 8/74 (10.81%)# |
| Sole deletion and/or duplication | 8/74 (10.81%) | 11/50 (22.00%) | 25/130 (19.23%) | 14/50 (28.00%) | 18/74 (24.32%) |
| Sole deletion and/or duplication euploidy embryos with mosaic forms | 3/74 (4.05%) | 9/50 (18.00%) | 10/130 (7.69%) | 7/50 (14.00%) | 9/74 (12.16%) |
| Sole aneuploidy | 7/74 (9.46%) | 5/50 (10.00%) | 20/130 (15.38%) | 4/50 (8.00%) | 14/74 (18.92%) |
| Aneuploidy embryos with mosaic forms | 4/74 (5.41%) | 0 | 6/130 (4.62%) | 3/50 (6.00%) | 3/74 (4.05%) |
| Aneuploidy embryos with deletion and/or duplication | 1/74 (1.35%) | 0 | 4/130 (3.08%) | 0 | 4/74 (5.41%) |
| Aneuploidy embryos with chromosomal deletion and/or duplication and mosaic forms | 2/74 (2.7%) | 1/50 (2%) | 5/130 (3.85%) | 1/50 (2%) | 2/74 (2.7%) |

*40.54% vs. 21.62%, OR = 2.47 (1.20–5.09), $P = 0.020$;

#25.68% vs. 10.81%, OR = 2.85 (1.16–7.01), $P = 0.032$;

Table 8. PGT-SR (errors from the proband) results by female PN 14 h coefficient β1 ranking.

| PGT-SR result | First 1 | First 2 | Median | Last 2 | Last 1 |
|---|---|---|---|---|---|
| Chromosomal normal embryo | 26/42 (61.9%)* | 10/27 (37.04%) | 32/73 (43.84%) | 12/27 (44.44%) | 10/42 (23.81%)* |
| Embryo's chromosomal error coincident with female | 12/42 (28.57%) | 13/27 (48.15%) | 28/73 (38.36%) | 10/27 (37.04%) | 21/42 (50%) |
| Embryo's chromosomal error not coincident with female | 4/42 (9.52%) | 4/27 (14.81%) | 13/73 (17.81%) | 5/27 (18.52%) | 11/42 (26.19%) |

*61.9% vs. 23.81%, OR = 5.20 (2.02–13.37), $P < 0.001$

Table 9. PGT-SR results (errors from the proband) and male PN 14 h coefficient β1 ranking.

| PGT-SR result | First 1 | First 2 | Median | Last 2 | Last 1 |
|---|---|---|---|---|---|
| Chromosomal normal embryo | 19/32 (59.38%) | 13/23 (56.52%) | 28/57 (49.12%) | 12/23 (52.17%) | 16/32 (50%) |
| Embryo's chromosomal error coincident with male | 9/32 (59.38%) | 7/23 (30.43%) | 23/57 (40.35%) | 6/23 (26.09%) | 6/32 (18.75%) |
| Embryo's chromosomal error not coincident with male | 4/32 (12.5%) | 3/23 (13.04%) | 6/57 (10.53%) | 5/23 (21.74%) | 10/32 (31.25%) |

Table 10. The female PN and total DNA content.

| Rank | 0 | DNA increase | decrease | |
|---|---|---|---|---|
| 1 | 89/155 (57.42%) | 32/155 (20.64%) | 33/155 (21.29%) | 155 |
| 2 | 79/155 (50.97%) | 40/155 (25.81%) | 36/155 (23.23%) | 155 |
| 3 | 78/138 (56.52%) | 39/138 (28.26%) | 21/138 (15.22%) | 138 |
| 4 | 46/107 (42.99%) | 25/107 (23.36%) | 36/107 (33.64%) | 107 |
| 5 | 40/81 (49.38%) | 21/81 (25.93%) | 20/81 (24.69%) | 81 |
| 6 | 33/56 (58.93%) | 12/56 (21.43%) | 11/56 (19.64%) | 56 |
| 7 | 23/37 (62.16%) | 9/37 (24.32%) | 5/37 (13.51%) | 37 |
| 8 | 13/25 (52.00%) | 8/25 (32.00%) | 4/25 (16.00%) | 25 |
| 9 | 8/13 (61.54%) | 4/13 (30.77%) | 1/13 (7.69%) | 13 |
| 10 | 6/12 (50.00%) | 3/12 (25.00%) | 3/12 (25.00%) | 12 |
| >10 | 2/11 (18.18%) | 5/11 (45.45%) | 4/11 (36.36%) | 11 |
| | 417 | 199 | 174 | 790 |

Correlation analysis: $P = 0.491$

Supplement

Fig. 1. The heterogeneity of female pronucleus area coefficient β1

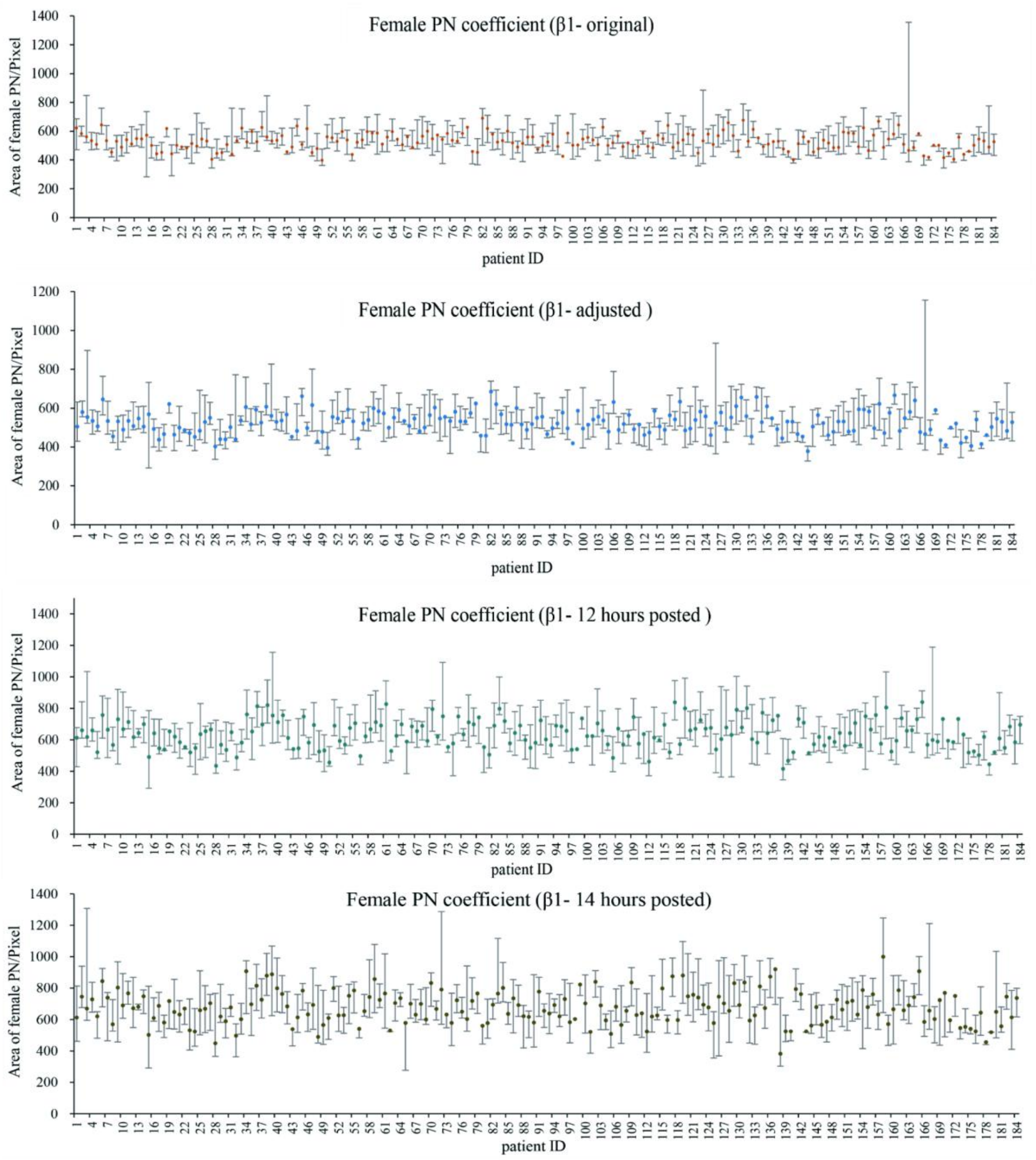

Supplement

Fig. 2. The definition of ranking by PN coefficient β1.

| Number of embryo | Ranking by PN coefficient β1 | | | | | | | | | | |
|---|---|---|---|---|---|---|---|---|---|---|---|
| 2 | 1 | 2 | | | | | | | | | First 1 |
| 3 | 1 | 2 | 3 | | | | | | | | First 2 |
| 4 | 1 | 2 | 3 | 4 | | | | | | | Medi |
| 5 | 1 | 2 | 3 | 4 | 5 | | | | | | Last 2 |
| 6 | 1 | 2 | 3 | 4 | 5 | 6 | | | | | Last 1 |
| 7 | 1 | 2 | 3 | 4 | 5 | 6 | 7 | | | | |
| 8 | 1 | 2 | 3 | 4 | 5 | 6 | 7 | 8 | | | |
| 9 | 1 | 2 | 3 | 4 | 5 | 6 | 7 | 8 | 9 | | |
| 10 | 1 | 2 | 3 | 4 | 5 | 6 | 7 | 8 | 9 | 10 | |
| 11 | 1 | 2 | 3 | 4 | 5 | 6 | 7 | 8 | 9 | 10 | 11 |

Fig. 3. Chromosome-normal rate in a female age-dependent distribution.

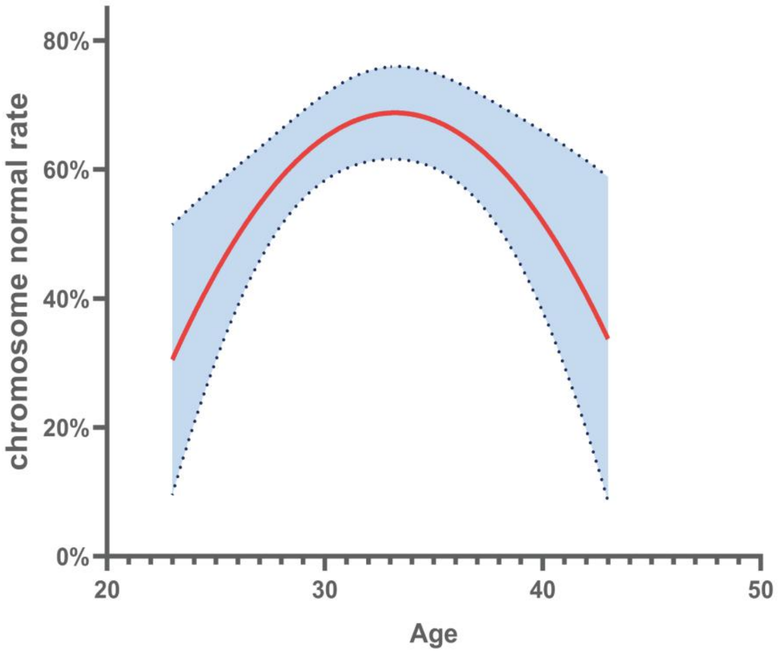
